# WWOX promotes osteosarcoma development via upregulation of Myc

**DOI:** 10.1101/2023.03.14.532523

**Authors:** Rania Akkawi, Osama Hidmi, Ameen Haji Yehya, Jonathon Monin, Judith Diment, Yotam Drier, Gary S. Stein, Rami I. Aqeilan

## Abstract

Osteosarcoma is an aggressive bone tumor that primarily affects children and adolescents. This malignancy is highly aggressive, associated with poor clinical outcomes, and primarily metastasizes to the lungs. Due to its rarity and biological heterogeneity, limited studies on its molecular basis exist, hindering the development of effective therapies. The WW domain-containing oxidoreductase (WWOX) is frequently altered in human osteosarcoma. Combined deletion of *Wwox* and *Trp53* using *Osterix1-Cre* transgenic mice has been shown to accelerate osteosarcoma development. In this study, we generated a traceable osteosarcoma mouse model harboring the deletion of Trp53 alone (single-knockout) or combined deletion of Wwox/Trp53 (double-knockout) and expressing a tdTomato reporter. By tracking Tomato expression at different time points, we detected the early presence of tdTomato-positive cells in the bone marrow mesenchymal stem cells of non-osteosarcoma-bearing mice (young BM). We found that double-knockout young BM cells, but not single-knockout young BM cells, exhibited tumorigenic traits both in vitro and in vivo. Molecular and cellular characterization of these double-knockout young BM cells revealed their resemblance to osteosarcoma tumor cells. Interestingly, one of the observed significant transcriptomic changes in double-knockout young BM cells was the upregulation of Myc and its target genes compared to single-knockout young BM cells. Intriguingly, Myc-chromatin immunoprecipitation sequencing revealed its increased enrichment on Myc targets, which were upregulated in double-knockout young BM cells. Restoration of WWOX in double-knockout young BM cells reduced Myc protein levels. As a prototype target, we demonstrated the upregulation of MCM7, a known Myc target, in double-knockout young BM relative to single-knockout young BM cells. Inhibition of MCM7 expression using simvastatin resulted in reduced proliferation and tumor cell growth of double-knockout young BM cells. Our findings reveal BM mesenchymal stem cells as a platform to study osteosarcoma and Myc and its targets as WWOX effectors and early molecular events during osteosarcomagenesis.

## Introduction

Osteosarcoma (OS) is the most common primary malignant bone tumor that predominantly affects children and adolescents. It often arises in the metaphysis of long bones(1). OS is highly malignant, with irregular bone growth and distant metastases that are commonly observed in the lungs. The current standards of care for OS are neoadjuvant chemotherapy, surgical resection, and adjuvant chemotherapy. This treatment increases the disease free-survival probability to more than 60% in the localized OS cases(2), but it is still between 10%-40% in metastatic OS(3). Molecular studies of OS are significantly hindered by its genetic complexity and chromosomal instability (4),(5). In addition, the absence or uncertainty of the exact cell of origin further challenges the identification of therapeutic targets(6). Therefore, an in-depth molecular understanding of osteosarcomagenesis is urgently needed for better prognosis and management of patients with OS.

WW-domain containing oxidoreductase (*WWOX*) spans one of the most active common fragile sites FRA16D(7), and encodes a 46kDa tumor suppressor that is altered in most human cancers(8),(9). We have previously demonstrated that WWOX is frequently altered in human OS and that WWOX restoration in WWOX-negative OS cells suppresses tumorigenicity(10). Furthermore, our group has shown that ablation of murine *Wwox* using *Osterix1(Osx)-Cre* transgenic mice with a p53-deficient background (DKO) accelerates OS development. DKO tumors display a more aggressive and poorly differentiated phenotype than p53 single KO (SKO) tumors. The DKO model highly recapitulates human OS and represents a valuable platform for investigating OS genetics and preclinical therapeutic targets(11).

Myc is a transcription factor encoded by the c-Myc gene which regulates an estimated 15% of genes in the human genome(12). Elevated Myc expression can be found in up to 70% of human tumors, and suppression of Myc is thought to lead to tumor regression(13). In OS, Myc is known to play a key role(14),(15), though the kinetic of its deregulation is not well defined. Myc may contribute to tumorigenesis by several ways including overstimulating cell growth and metabolism and/or by causing genomic instability(16), probably through its ability to induce DNA damage, promote gross chromosomal rearrangements, induce inappropriate cell cycle progression and impair DNA repair(17). Moreover, Myc has been reported to regulate the pre-replicative complex of the cell cycle including MCM5 and MCM7 among other proteins(18),(19). The minichromosome maintenance proteins (MCMs), which exist in a functional complex consisting of MCM2-7 play important roles in double-strand DNA unwinding, DNA replication control, and DNA damage repair. Various cancer tissues, including sarcoma and carcinoma cell lines, show overexpression of MCMs, particularly MCM7 (20),(21),(22). Inhibition of MCM7 results in the accumulation of DNA lesions and the activation of the DNA damage checkpoint response hence preventing cell growth(23). Interestingly, MCM7 has been shown to be targeted by statins in Rb-deficient tumors(24). In addition, simvastatin has been shown to suppress tamoxifen-resistant cell growth by inhibiting MCM7 and Rb and subsequently inducing DNA damage(25). However, the role of statins in the inhibition of MCM7 in OS is unknown.

Mesenchymal stem cells (MSCs) or osteoblasts can be the cells of origin for OS, although this remains controversial. Research evidence suggests that OS originates during the differentiation process of MSCs into preosteoblast, and that the stage of differentiation affects the OS phenotype and grade(26),(27). In contrast, others suggest that osteoblast precursor cells are the cells of origin for OS(6),(28). Recently, it was shown that *Osx* is expressed in both the osteoblast lineage (osteoblasts, osteocytes, and chondrocytes) and in bone marrow (BM) stromal cells, BM-adipocytes, and BM-perivascular cells(29). This finding prompted us to use our SKO and DKO models to dissect the early molecular changes in OS development. To this end, we introduced a tdTomato reporter into these models and followed their behavior at different time points. We found that tdTomato-expressing BM isolated from young DKO mice (DKO yBM), prior to OS development, can form OS tumors that are indistinguishable from DKO tumors. This feature was not observed in the tdTomato-expressing BM isolated from young SKO mice (SKO yBM). Molecular characterization of DKO yBM cells revealed unique upregulation of several oncogenic pathways including upregulation of Myc and its targets at very early stages prior to OS onset. Inhibition of MCM7, a known Myc target, attenuates OS tumor formation. Our model represents a unique platform for studying early changes and molecular targets during osteosarcomagenesis and for assessing vulnerabilities that can be targeted in OS.

## Materials and Methods

### Mouse strains and tumor models

Mice were housed in a pathogen-free animal facility at the Hebrew University of Jerusalem. To generate our traceable mouse model, we crossed the single knockout (*Trp53*) or double knockout mice (*Wwox*/*Trp53*) Osx1-Cre for deletion of *Trp53* or *Trp53*/*Wwox* respectively in mesenchymal progenitors committed to bone formation(11) with B6.Cg-Gt(ROSA)26SorTm9(CAG-tdTomato) Hze/J (TdT) (catalog #007909; The Jackson Laboratory) to generate SKO-Osx-Cre;tdTomato (SKO^TOM^ or in short SKO) and DKO-Osx-Cre;tdTomato (DKO^TOM^ or in short DKO). Bone Marrow (BM) and tumor cells were collected from mice at different time points.

For Intratibial Injection (IT), 6-8 weeks old NOD/SCID mice were anesthetized with ketamine (80 mg/kg) (Vetoquinol) / xylazine (7 mg/kg) (Phibro) and injected intraperitoneally with 5 mg/kg carprofen (Norbrook). Mice were then injected intratibially with 0.5×10^6^ tumor or BM cultured cells (as explained in BM extraction and cell culture section), suspended in 20μl sterile complete media (MEM-Alpha (Biological Industries)) supplemented with 10% Fetal Calf Serum (FCS) (Gibco) and 1% L-Glutamine/Penicillin/Streptomycin (Biological Industries). 12-hours post IT-injection, the mice were administered with another dose of carprofen (5 mg/kg). The mice were monitored for about one-four months for tumor development.

For intravenous injection, a total of 1×10^6^ cells suspended in complete MEM-α, were injected in the tail vein of 6-8 weeks old NOD/SCID mice to study the colonization ability of cells in lungs. After 3-4 months postinjection, mice were sacrificed, and lungs were collected for further analysis(30).

To validate the inhibitory effect of simvastatin (SVA) *in vivo*, NOD/SCID mice that developed OS tumors after IT injection were split randomly into two groups, mice that received 60 mg/kg/day SVA (Sigma) (8-mice), and a vehicle group (6-mice). The two groups received a gavage injection for 10 days, then the mice were euthanized, and tumors were collected for further analysis(24).

#### Ethical Compliance

All procedures performed in this study involving animal use were in accordance with the ethical standards of the institutional and/or national research committee.

### Bone Marrow (BM) extraction and culture

BM cells were collected from WT mice (control), young non-tumor-bearing SKO and DKO-Osx1-Cre-tdTomato mice at different time points (SKO yBM, DKO yBM respectively), as well as at the time of tumor detection (SKO BMT, DKO BMT respectively). Mice were sacrificed using CO_2_, femurs and tibiae from both legs were cleaned from connective tissue, and the BM was flushed out using a syringe with 27 Gauge needle. Red blood cells were lysed using a hypotonic treatment (water dilution). The cells were then centrifuged at 2000 rpm for 10 min, filtered using 70µM cell strainer (Corning, Massachusetts, USA) and plated at an appropriate density after one passage (p1)(31).

### Tumor dissection and tumor cell lines preparation

At the time of tumor detection, mice were euthanized, and tumors were extracted and minced using a sterile scalpel and digested with trypsin-EDTA (25 mg/ml) (Sigma), Collagenase P (1 mg/ml) (Sigma Cat# 11213865001), and DNase I (10 mg/ml) (Sigma Cat# 10104159001) for 1hour at 37°C with vigorous shaking. Cells were then filtered through 70µM cell strainer, centrifuged at 2000 rpm for 10 min, and plated in 10 cm^2^ plates in MEM-α medium supplemented with 10% FCS and 1% L-Glutamine/Penicillin/Streptomycin.

### Flow Cytometry analysis of the mesenchymal and hematopoietic markers

1×10^6^ of yBM, BMT, and tumor cells were collected freshly (direct) and/or after culture (Cult)-when the tdTomato positive cells were dominant. Cells were stained with FACS buffer (2% FCS, 1mM EDTA, 0.1% NAN_3_ in PBS), containing the following antibodies: for mesenchymal markers, Sca-1-APC (clone D7, Cat# 108111, Biolegend,1:250), CD29-Alexa Flour 700 (clone HMβ1-1, Cat# 102218, BioLegend, 1:250), and for hematopoietic markers; CD45.2-Brilliant Violet 421 (clone 104, Cat# 109831, BioLegend, 1:100), for 30 min at 4 °C. Samples were acquired using a CytoFlex cytometer and analyzed using CytExpert version 2.4(32).

### Osteogenic Differentiation of yBM, BMT and tumor cells

yBM, BMT, and tumor cells were induced to differentiate by plating them at 70% confluency, and culturing in osteogenic medium that included MEMα supplemented with 10% FCS, Ascorbic Acid (50μg/ml) and β-glycerophosphate (3mM). Osteogenic medium was added and replaced every 2-3 days for 21 days. Mineralization was assessed by measuring the expression levels of osteogenic markers (real-time PCR-as explained later in real time PCR section), and the formation of calcium-enriched nodules was detected by Alizarin Red Staining(11).

### Colony formation assay

Five hundred yBM, BMT, and tumor cells were seeded in 6-well plates (triplicate), and the colonies were monitored for one week. The colonies were then fixed using 70% ethanol for 10 min and stained with Giemsa stain for 30 min. The colonies were then washed and counted(33) The graph shows fold change/number of colonies compared to the control groups, and bars represent standard deviation (SD).

To determine the effect of simvastatin (SVA) on the survival of yBM, BMT, and tumor cells, 24 h after seeding, SVA was added at the indicated concentrations for 48 h, and the colonies were monitored for one week. Fixation, staining, and analysis were performed as previously described.

### Wound Healing-Scratch Assay-Migration

The migration abilities of yBM, BMT, and tumor cells were determined using a wound healing scratch assay. 3×10^4^ yBM, BMT, and tumor cells were plated onto an IncuCyte 96-Well ImageLock plate (IncuCyte 4379) for 12 h. Then, the confluent monolayer was washed with PBS and a scratch was made using IncuCyte: 96-pin wound-making tool (IncuCyte WoundMaker), and then the wells were washed three times with culture media. After that, serum-free media (SFM) was added. The plates were incubated in IncuCyte live-cell imaging system, and the 96-well wound assay protocol was run on the software. yBM, BMT and tumor cells were imaged at 2 h intervals for 72h. Analysis was performed using the Cell Migration Analysis software (IncuCyte 4400)(34).

### Wound Healing-Scratch Assay-Invasion

The invasive abilities of yBM, BMT, and tumor cells were determined using a wound healing scratch assay. An IncuCyte 96-Well ImageLock plate was coated with Matrigel (BD 354230, 1:20 with serum-free media) for 12 h. Next, 3×10^4^ yBM, BMT, and tumor cells were plated onto an IncuCyte 96-Well ImageLock plate (IncuCyte 4379) for 12 h. Then, the confluent monolayer was washed with PBS and a scratch was made using IncuCyte: 96-pin wound-making tool (IncuCyte WoundMaker), and then the wells were washed three times with culture media. After that, 100µl of media was added. The plate was cooled to 4 °C for 5 min, then the media was removed, a top layer of Matrigel (50µl) was added, and the plate was incubated at 37°C for 30 min, before adding another 100µl of serum-free media. The plates were incubated in IncuCyte live-cell imaging system, and the 96-well wound assay protocol was run on the software. yBM, BMT and tumor cells were imaged at 2 h intervals for 72h. Analysis was performed using the Cell Migration Analysis software (IncuCyte 4400)

### Cell Proliferation assay (MTT assay)

To check the effect of simvastatin (SVA) on cell viability, yBM, BMT, and tumor cells (3×10^3^) were plated onto 96 well plate (triplicate for each group and for each concentration), at 24h intervals. Then, activated SVA (as explained later) was added at the indicated concentrations for 48 h and analyzed using the MTT reagent according to the manufacturer’s instructions (Promega). The graph represents the comparison with the control group, and the bars represent ± the SD.

For SVA activation, 4 mg of simvastatin was dissolved in 100µl of ethanol. Then 150µl of 0.1NaOH was add to the solution and incubated at 50°C for 2 h. The pH was brought to 7.0, and the final concentration of the stock solution was 4 mg/ml. The stock solution was stored at 4 °C(35).

### Plasmid-WWOX manipulation

For generating WWOX overexpression in DKO yBM cells, cells were infected with lentiviral vector containing the wild type WWOX and selected with 2 mg/mL G418 (Gibco: 11811031)(36)

### Ionizing radiation

Cre+ WT BM, SKO, DKO yBM cells were irradiated with 10Gy, then collected at several time points (30, 120 min) after irradiation(37)

### Immunoblotting

Total protein was lysed by using lysis buffer containing 50 mM Tris (pH7.5), 150 mM NaCl, 10% glycerol, 0.5% Nonidet P-40 (NP-40), with addition of protease and phosphatase inhibitors (1:100). The lysates were subjected to SDS-PAGE under standard conditions. The following antibodies were used: rabbit polyclonal WWOX 1:10000, mouse monoclonal Gapdh (CB-1001-), Calbiochem 1:10000, mouse monoclonal p21(F-5), sc 6246, Santa Cruz 1:200, mouse monoclonal p53 (1C12), Cell Signaling, 1:1000, mouse monoclonal MCM7 (141.2), sc-9966-Santa Cruz 1:200, rabbit polyclonal c-Myc (9402)-Cell Signaling, 1:750, rabbit polyclonal DsRed (632496), TaKaRa, 1:500, and mouse monoclonal β-actin (A5441), Sigma 1:10000.

### Histology and Immunohistochemistry

Fixed bones and tumors [4% formalin] were partially decalcified in 14% EDTA for three days, followed by paraffin embedding, sectioning then staining with hematoxylin (Sigma)) and eosin (Sigma) (H&E). For Immunohistochemistry, paraffin-embedded tissues were deparaffinized, followed by antigen retrieval with 25mM citrate buffer (PH 6) in a pressure cooker. Then, the sections were left to cool for 25 min, followed by blocking of the endogenous peroxidase with 3% H_2_O_2_ (Sigma) for 15 min. To reduce non-specific binding of the primary antibody, tissues were blocked with blocking solution (CAS Block) (Invitrogen), followed by incubation with the primary antibody overnight at 4°C. Sections were washed with TBST, followed by incubation with secondary anti-mouse immunoglobulin antibody for 30 min (Nichirei Biosciences). The reaction was then performed using a DAB peroxidase kit (Vector Laboratories, SK-4100), followed by hematoxylin (Bar Naor, Cat# BN4537N) staining for 40s as a counterstain. IHC stains were manually counted, at least five pictures were taken for each slide, and the average was calculated for each slide. The antibodies used were mouse monoclonal MCM7 (141.2) [sc-9966-Santa Cruz 1:200].

### Immunofluorescence

Bones (Femur and Tibia) were fixed in 4% PFA overnight at 4 °C. Fixed bones were partially decalcified in 14% EDTA for 3 days, washed in a sucrose gradient (1 h in 10% sucrose, 1 h in 20% sucrose, overnight in 30% sucrose) before snap-freezing them in OCT embedding medium. Frozen sections were cut at 10μm thickness and stored at −80 °C. Sections were warmed to room temperature, washed in PBS, fixed with 4% PFA for 10 minutes then washed two times with PBS. The sections were incubated with Hoechst 33258 diluted in 5% normal goat serum, 0.05% BSA and 0.1 %Triton X-100 in PBS for 15-minutes. The slides were then dried and mounted with coverslips using immunofluorescence mounting medium (Dako s3023). Sections were then imaged using an Olympus FLUO-VIEW FV1000 confocal laser scanning microscope and processed using the associated Olympus FLUO VIEW software(38),(39).

For paraffin-embedded sections, after deparaffinization and antigen retrieval, as mentioned above, tissues were blocked using a blocking solution (5% normal goat serum, 0.05% BSA, and 0.1 %Triton X-100 in PBS) for 40-minutes. The sections were then incubated with the primary antibody diluted in the blocking solution overnight at 4°C °C. The following day, the sections were washed 3-times with PBS containing 0.05% tween-20 (PBST), incubated with secondary antibody and Hoechst 33258 diluted in blocking solution for 1-hour at room temperature, then washed 3-times with PBST. The slides were then dried, mounted, and imaged as described above. The antibody used was rabbit polyclonal DsRed (632496; TaKaRa, 1:500).

### Real Time PCR

Total RNA was extracted using TRI reagent (Biolab), as described by the manufacturer for the phenol/chloroform-based method. cDNA was prepared with 1μg of RNA using a QScript cDNA synthesis kit (Quantabio). The SYBR Green PCR Master Mix (Applied Biosystems) was used for qRT-PCR. All measurements were performed in triplicate and standardized to HPRT levels. The primer sequences are listed in Supplemental Table 1.

### RNA Sequencing and analysis

Four WT, six non-tumor bearing, and six tumor-bearing mice were euthanized by CO_2_, and BM and/or tumor were extracted and cultured as described above. After three passages, tdTomato-positive cells were dominant in the culture, and RNA was prepared using the TRI reagent (Biolab). Library preparation and sequencing were performed by the Genomic Applications Laboratory in Hebrew University’s Core Research Facility using a standard protocol. RNA quality was checked using a tape station (Agilent Technologies), followed by poly-A cleanup and cDNA preparation. To prepare cDNA, 1 µg of RNA from each sample was processed using the KAPA Stranded mRNA-Seq Kit with mRNA Capture Beads (Kapa Biosystems). Sequencing was performed using NextSeq 500 (Illumina). For library quality control, measures were applied to raw FASTQ files, trimming low-quality bases (Q>20), residual adapters, and filtering out short reads (L>20). These measurements were performed with the Trim-Galore (v0.4.1) wrapper tool. File assessment before and after quality control was performed using FastQC (v0.11.5). The mouse transcriptome was mapped using salmon v0.13.1. Differential gene expression for the 16 samples was explored using the R package (v1.16.1) and analyzed using DESeq2 (v1.28.1).

### Chromatin immunoprecipitation sequencing (ChIP-seq)

ChIp-seq was performed as previously described(40). Briefly, BM DKO cells (∼10^7^) were crosslinked with 1% formaldehyde (methanol free, Thermo Scientific 28906) for 10 min at room temperature and quenched with glycine, 125 mM final concentration. Fixed cells were washed twice in PBS and incubated in lysis buffer (10mM EDTA, 0.5% SDS, 50mM Tris-HCl pH=8, and protease and phosphatase inhibitors) for 30min on ice. Cells were sonicated using bioruptor sonicator to produce chromatin fragments of ∼200-300 bp. The sheared chromatin was centrifuged 10min at 20000g. From the supernatant, 2.5% were saved as input DNA and the rest was diluted in dilution buffer (50mM TRIS-HCl pH8, 0.01%SDS, 150mM NaCl, and 1% Triton X-100). The chromatin was immunoprecipitated by incubation with 5 μl of anti-c-Myc antibody (rabbit polyclonal c-Myc (9402)-Cell Signaling,). Immune complexes were captured with protein G Dynabeads. Immunoprecipitates were washed once with low salt, twice with high salt buffer, and twice with LiCl buffer, and twice with TE buffer. The chromatin was eluted from the beads with 240 μl of elution buffer (100mM sodium bicarbonate and 1% SDS) and incubated overnight at 65°C to reverse the cross-linking. Samples were treated with proteinase K at 45 °C for 2 h. DNA was precipitated by phenol/chloroform/isoamylalcohol extraction. The ChIPed and the Input DNA were used to prepare libraries by Kappa Hyperprep kit and sequenced in Nextseq (Illumina).

### Data Access Statement

Raw data are available upon reasonable request.

### Statistical analysis

The results of the experiments are expressed as mean± SD. The student’s t-test was used to compare the values of the test and control samples. p-value ≤ 0.05 indicates a significant difference; ** p ≤ 0.01, *** p ≤ 0.001, **** p ≤ 0.0001.

## Results

### Generation of a traceable mouse model of osteosarcoma

We have recently shown that depletion of both *Wwox* and *p53* (DKO) in osteoblast progenitors, using *Osterix1-Cre*, accelerates osteosarcoma (OS) development, resulting in more aggressive and chemotherapy-resistant OS than *p53* knockout alone (SKO)(11). To better dissect the early molecular events contributing to osteosarcomagenesis, we bred our DKO mouse model with *Rosa26-LSL-tdTomato* transgenic mice (Figure 1A). Depletion of *Wwox* and *p53* and the insertion of *tdTomato* (DKO^TOM^) were confirmed by genomic PCR (Figure 1B). Expression of the tdTomato reporter in Osterix-expressing cells was validated by fluorescence microscopy (Figure 1C). Consistent with previous work(29), tdTomato-positive cells expressing *Osterix* were also detected in the bone marrow (BM) of young non-tumor-bearing mice, as validated by immunofluorescence of bone-frozen sections (Figure 1D) and flow cytometry analysis of BM cells representing the percentage of tomato positive cells in the BM of 2-months old mice (10.2%) (Figure1E). Ablation of WWOX and p53 in cultured tdTomato-positive BM cells was also confirmed at the protein level after exposure to ionizing radiation (10 Gy) by immunoblot analysis (Figure 1F). In agreement with our previous findings, DKO^TOM^ mice developed OS with a latency of 8-months (data not shown), as previously shown for DKO(11). Using this new mouse model, we next determined the molecular and cellular changes that occur during OS development.

**Figure 1:**
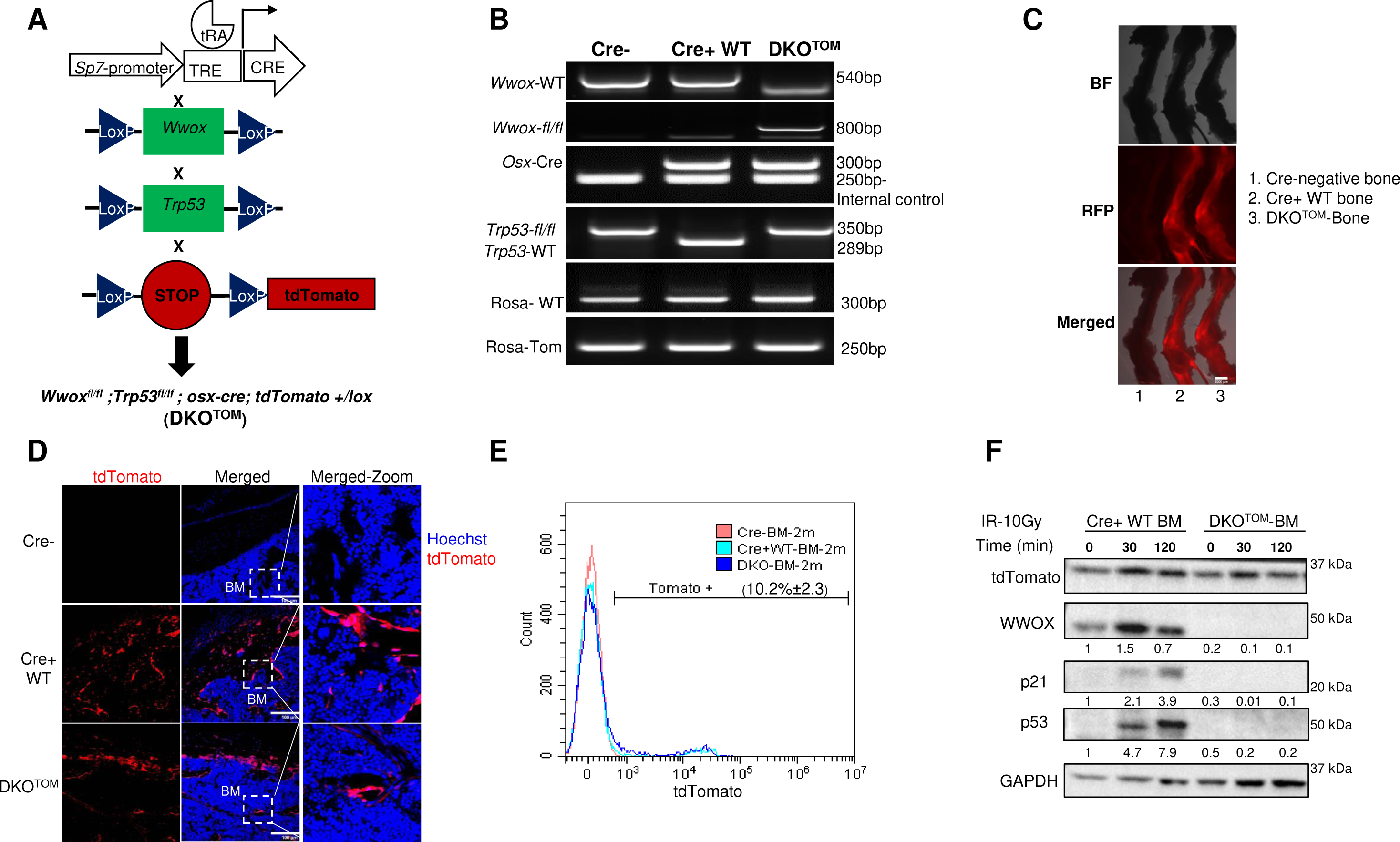
Generation of a traceable mouse model of osteosarcoma cells. **(A)** Schematic illustration of the experimental OS mouse model. **(B)** Genomic PCR of Cre negative (Cre-), Cre positive (Cre+ WT) and Double knock out (DKO^TOM^) mice. Labels in the left panel represent the primer pairs and in the right panel represent the expected band size. **(C)** Fluorescent microscope images for bones extracted from (1) Cre-negative mouse, (2) Cre+WT mouse, (3) DKO^TOM^ mouse. **(D)** Immunofluorescence images of frozen bone sections from Cre-, Cre+WT, and DKO^TOM^ mice. Red signal represents the endogenous tdTomato, blue signal represents the nuclear staining-hoechst. **(E)** Flow cytometry analysis of BM cells of 2-months (2m) old Cre-, Cre+WT, and DKO^TOM^ mice. 10.2±2.3 represents the percentage of tdTomato positive cells in the Cre+ WT and DKO BM compared to the Cre-mice of similar age. **(F)** Immunoblotting of p53, WWOX, p21, and tdTomato in the Cre+ WT and DKO BM cells before (0 min) and after (30, 120 min) of ionizing radiation exposure (IR-10Gy). (Quantification relative to Cre+ WT is represented under the blot). **WT**: Wildtype, **DKO**: Double Knock Out, **BF**: Bright Field, **RFP**: Red Fluorescent Protein, **BM**: Bone Marrow.

### Bone marrow cells of OS-bearing mice are tumorigenic

To elucidate the kinetics of OS tumor formation in DKO^TOM^ mice, we examined the presence of tomato florescence in different compartments. Interestingly, we detected a high percentage of tdTomato-positive cells both in the directly collected tumor (before culturing) [Direct-Tumor] and BM of DKO^TOM^ tumor-bearing mice (before culturing) [Direct-BMT] (Figure 2A). These results prompted us to test whether BMT cells exhibit tumorigenic abilities. To this end, we isolated and cultured tdTomato BMT and tumor cells and examined their ability to survive and proliferate *in vitro*. We found that BMT cells were able to form colonies similar to those isolated from the DKO^TOM^ tumor itself (Figure 2B), as compared to BM cells isolated from wild-type mice (Cre+WT;tomato), which failed to form any colonies, as expected. We next examined the tumorigenic capability of BMT cells *in vivo* and found that 78% of the immunocompromised mice that were intratibially (IT) injected with BMT cells were able to develop tomato-positive osteoblastic OS, as validated by H&E and immunofluorescence staining, respectively (Figure 2C). As expected, mice injected with wild-type BM cells (control) did not form any tumors. Notably, it took more time for BMT cells to develop tumors as compared to the paired primary tumor cells, suggesting that other hits are earned in the tumor itself (Figure 2D). More interestingly, these BMT cells exhibited pro-metastatic properties, as assessed by wound healing ability *in vitro*, both migration and invasion (Figure 2E, F).

**Figure 2:**
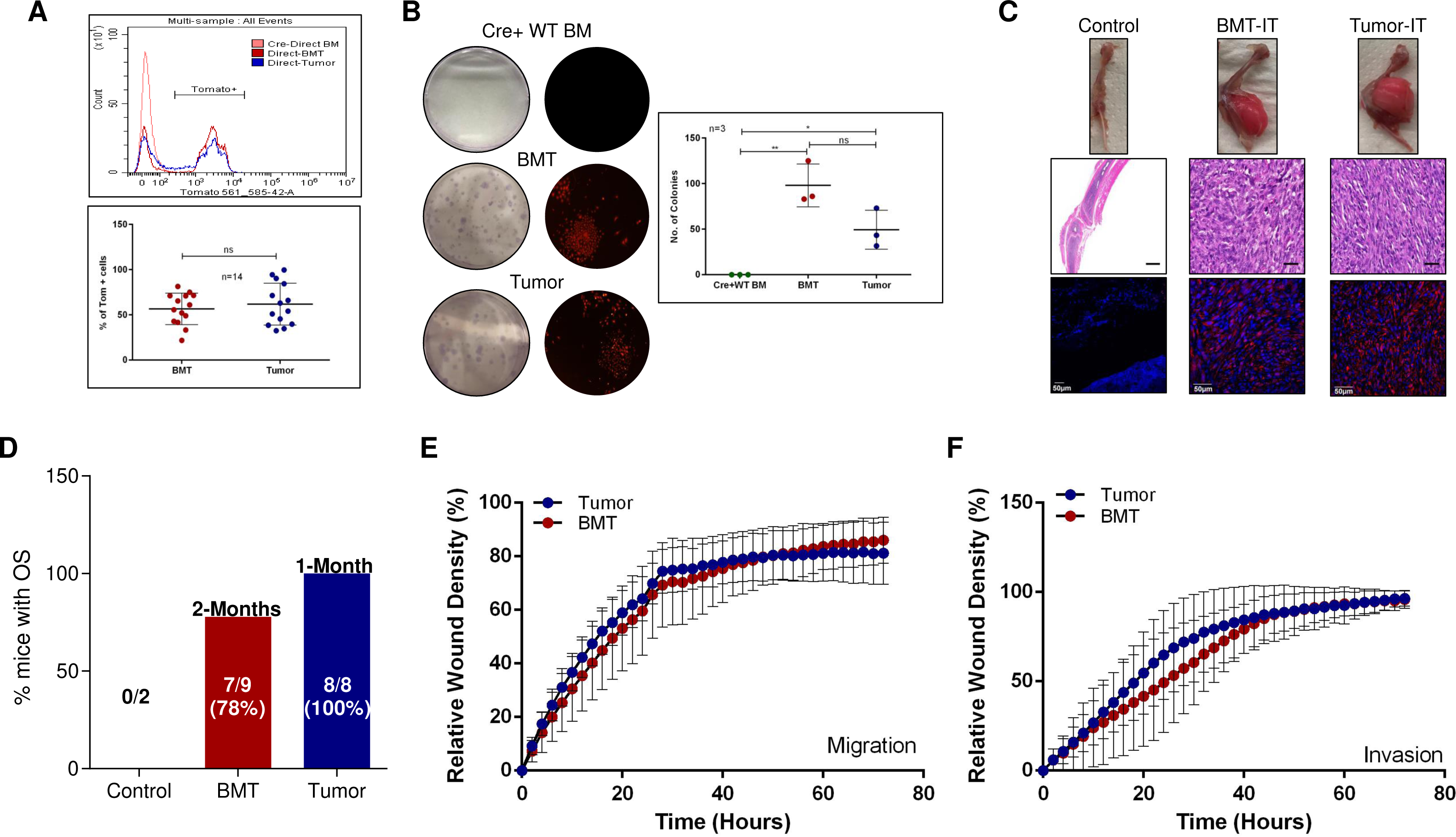
BM cells of OS-bearing mice are tumorigenic and exhibit metastatic ability. **(A)** Flow cytometry analysis of directly collected BM from a Cre-mouse (Cre-Direct BM), tumor (Direct-Tumor) and paired BM cells (Direct-BMT) collected from DKO^TOM^mouse demonstrating the percentage of tdTomato expression (representative experiment-upper panel, average of the percentage of tdTomato positive cells from 14 mice-lower panel). **(B)** Colony formation assay of paired BMT and tumor cells compared to Cre+WT BM (representative images-left panel (Giemsa stain and fluorescent microscope images), quantification of colony numbers of Cre+ BM, BMT and tumor cells-right panel (n=3). **(C)** Representative images of the OS tumors developed in NOD/SCID mice after intratibial (IT) injection of BMT and tumor cells compared to the control-upper panel, H&E validating the histological characteristics of OS-middle panel. Histological validation reveals OS of osteoblastic type. Immunofluorescence images of the developed tumors-lower panel. Red signal represents the endogenous tdTomato, blue signal represents the nuclear staining-hoechst. **(D)** Bar graph representing the percentage of immunocompromised mice that developed OS after IT injection of control BM cells (Control), BMT and tumor cells. **(E)** Wound healing assay-migration. Time plot representing the Relative Wound Density of tumor and BMT cells (n=3). **(F)** Wound healing assay-invasion. Time plot representing the Relative Wound Density of tumor and BMT cells (n=3).

Next, we sought to characterize the identity of the BMT cells. Immunophenotyping of the directly isolated BMT cells [direct]; before culturing; revealed a high percentage of cells expressing the hematopoietic cell surface marker (CD45.2) and the presence of a minor population of mesenchymal origin, expressing CD29 and Sca-1-mesenchymal cell surface markers (Figure 1S-A, Bi), consistent with a recent study confirming the expression of osterix in the hematopoietic lineage(41). Similar results were obtained for the tumor cells (Figure 1S-A, Bii). However, culturing of BMT and tumor cells [culture] led to the expansion of the mesenchymal population that was tdTomato-positive and diminished the hematopoietic population (Figure 1S-A, Bi-Bii). Importantly, these cultured BMT and tumor-tdTomato-positive cells expressed high levels of osteogenic markers, indicating their osteoblast identity (Figure 1S-C). In addition, BMT cells were able to differentiate toward the osteoblastic lineage, as validated by Alizarin red staining and mRNA levels of osteoblast markers, although less efficiently than tumor cells. (Figure 1S-D, E).

After demonstrating the tumorigenic ability of BMT cells, we investigated the molecular characteristics contributing to OS development. Therefore, we performed RNA sequencing (RNA-seq) analysis on the cultured control BM, BMT, and primary OS tumor cells of DKO mice. Using principal component analysis (PCA), we found that BMT and tumor cells clustered together in PC2 away from the control BM cells (Figure 3A). Strikingly, these BMT cells had 164 mutual differentially expressed (DE) genes in the tumor cells, as shown in the Venn diagram analysis (Figure 3B). Using unsupervised clustering analysis, we found that many OS-known genes were differentially upregulated (*Rrp9, Rcc1, Ran*), while others were downregulated (*Mylk, Nid2*) in the BMT and tumor cells compared to the control BM cells (Figure 3C). Pathway enrichment analysis and gene set enrichment analysis (GSEA) revealed significant changes in genes associated with the cell cycle (E2F and Myc targets) and DNA repair pathways (Figure 3D, E). Taken together, these data suggest that BMT cells isolated from OS-bearing mice are tumorigenic and, to a large extent, mirror OS tumor cells.

**Figure 3:**
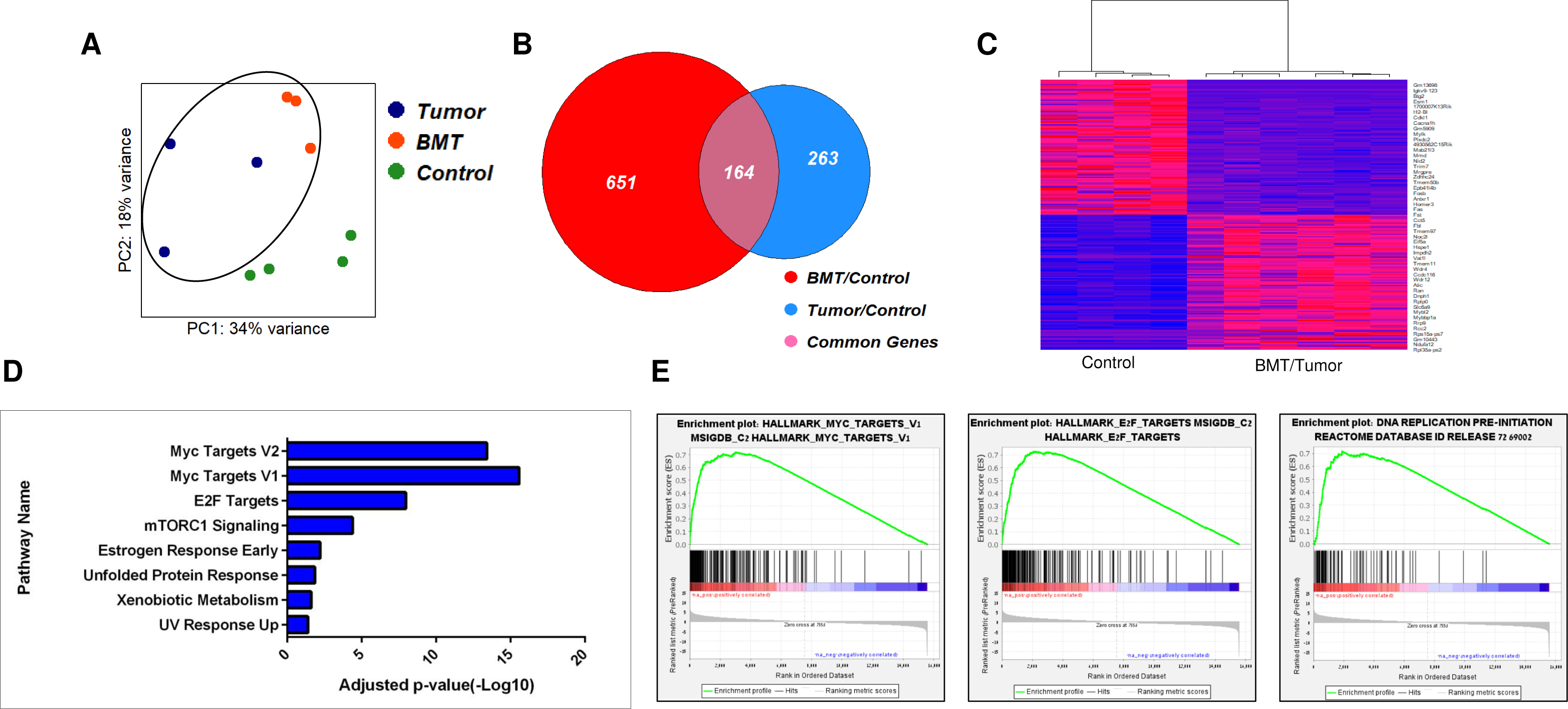
Molecular similarity between BMT and tumor cells. **(A)** Principal component analysis (PCA) plot of tumor cells (n=3), their paired BM cells (BMT) (n=3) and control BM cells (n=4). (B) Venn diagram analysis of tumor cells (n=3), their paired BM cells (BMT) (n=3) compared to the control. 164 are common genes between BMT and tumor cells. **(C)** Heat map illustration showing differentially expressed genes in Control and BMT/Tumor cells. **(D)** Pathway enrichment analysis **(E)** Gene Set Enrichment Analysis (GSEA) of differentially expressed genes between Control and BMT/tumor cells.

### young BM (yBM) cells exhibit tumorigenic ability

Our data imply that the BMT cells of DKO mice have tumorigenic abilities. Hence, we hypothesized that they might be used as proxies to dissect early molecular events contributing to osteosarcomagenesis. To address this, we collected BM cells from young non-tumor bearing mice (yBM) of different ages (1.5-6 months) of DKO mice and examined their cellular and molecular features. Intriguingly, we found that there was an increase in the percentage of tdTomato-positive cells in both wild type and genetically manipulated (DKO) yBM cells, which expanded with time in the latter (Figure 4A).

**Figure 4:**
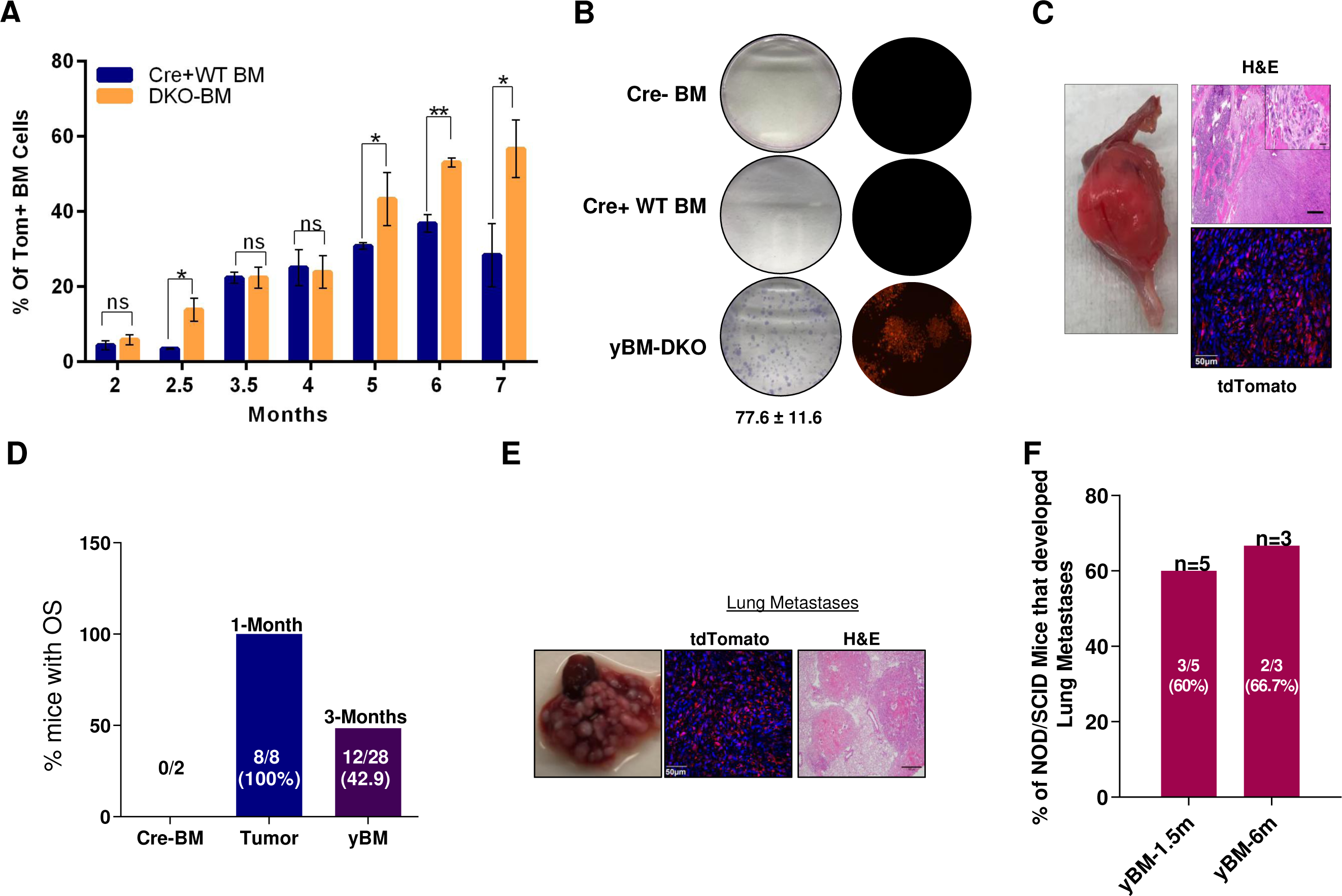
young BM (yBM)-derived cells exhibit tumorigenic and metastatic ability. **(A)** Summary graph of flow cytometry analysis of the percentage of tdTomato positive cells in the genetically manipulated BM (DKO BM) and wild type BM cells (Cre+WT BM) at different ages (n=3). **(B)** Colony formation assay of BM cells from young non-tumor bearing mice (yBM-DKO) (n=9), BM cells from Cre negative mice (Cre-BM) (n=3) and Cre+ WT mice (Cre+ WT BM) (n=3). Giemsa-stained colonies-left panel, fluorescent microscope images of the colonies-right panel, 77.6±11.6-represents the number of colonies in yBM-DKO. **(C)** OS tumors developed in immunocompromised mice after 3-4 months of Intratibial (IT)-injection of yBM cells. H&E staining of the developed OS tumors. Histological validation represents fibroblastic/osteoblastic OS-upper panel. Immunofluorescence staining of the developed tumors. Red signal represents the endogenous tdTomato, blue signal represents the nuclear staining-hoechst. **(D)** Bar graph representing the percentage of immunocompromised mice that developed OS following IT injection of yBM, tumor and control (Cre-BM). **(E)** Light microscope images, tdTomato immunofluorescence and H&E staining respectively representing lung metastasis in NOD/SCID mice after intravenous injection (IV) of yBM cells. **(F)** Bar graph representing the percentage of NOD/SCOD mice that developed lung metastasis after IV injection of yBM cells (collected from 1.5 and 6-months old mice).

We next validated that the yBM cells were of mesenchymal origin (Figure 2S-A, B), as observed previously for BMT cells. In addition, these yBM cells exhibited increased levels of osteogenic markers (Figure 2S-C) and differentiation capability, as assessed by Alizarin Red staining and qPCR analysis (F1S-D, E,). Remarkably, yBM cells isolated from DKO mice were able to form colonies (Figure 4B) and generate OS tumors upon IT injection in immunocompromised mice (Figure 4C). The latency of forming OS tumors was 3-months with penetrance that ranged from 36%-50% (Figure 4D). Histological analysis of these tomato-positive tumors revealed both fibroblastic and osteoblastic characteristics (Figure 4C). Altogether, these findings support our initial assumption that the BM cells in our DKO OS model can be used as a surrogate to investigate osteosarcomagenesis.

As the main hurdle in OS is the detection of lung metastasis at diagnosis, we tested whether these DKO yBM cells from non-tumor-bearing mice could exhibit migration and invasion abilities. Remarkably, DKO yBM cells, as well as BMT and tumor cells, demonstrated migration and invasion abilities, as validated by a wound healing assay *in vitro* (Figure 2S D, E). To further validate migration and invasion abilities *in vivo*, we intravenously injected yBM cells into NOD/SCD mice. Surprisingly, these cells were able to colonize the lungs and form nodules, as validated by immunofluorescence and H&E staining (Figure 4E), as indicated in the bar graph (Figure 4F). Taken together, these data suggest that DKO yBM cells isolated from non-OS-bearing mice display tumorigenic traits that could potentially reflect changes in OS tumor cells.

### Early molecular changes contributing to osteosarcomagenesis

Our previous results have demonstrated the tumorigenic ability of yBM cells isolated from non-OS-bearing mice. Next, we investigated the molecular characteristics of yBM DKO cells that contribute to OS development. Therefore, we performed RNA-seq analysis on control BM, yBM, and primary OS tumor cells of DKO mice. Using PCA, we found that the yBM DKO and tumor cells clustered together away from the control BM cells (Figure 5A). Strikingly, these yBM cells had 303 mutual genes with the tumor cells, as shown in the Venn diagram (Figure 5B). In addition, when we compared the DE genes between yBM DKO and BMT cells, they clustered together away from the control BM cells (Figure 5D). As shown in Figure 5E, 471 genes were common between yBM and BMT cells. Using unsupervised clustering analysis, we found that the upregulated and downregulated DE genes in the yBM and tumor cells, and in the yBM and BMT cells clustered together away from the control BM cells (Figure 3S-A, B). Pathway enrichment analysis and GSEA between yBM and tumor cells and between yBM and BMT cells revealed significant changes in genes associated with the cell cycle (E2F targets, Myc targets, and G_1_S transition) and DNA repair pathways (Figure 5C, F, G, H), consistent with those observed between BMT and tumor cells (Figure 3).

**Figure 5:**
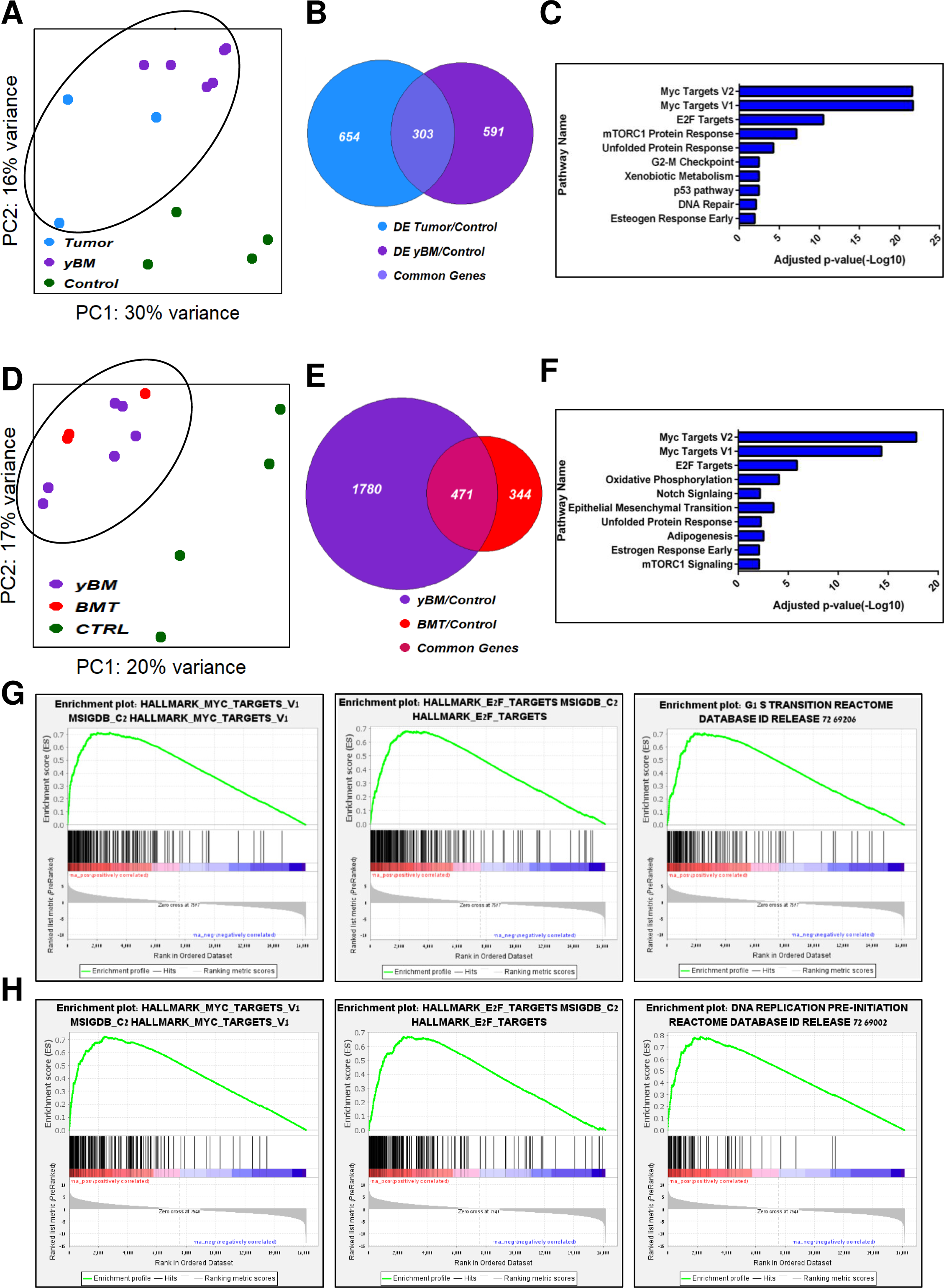
Early genes contributing to osteosarcomagenesis (yBM vs tumor cells and yBM vs BMT cells) **(A)** Principal component analysis (PCA) plot of tumor cells (n=3), yBM cells (1.5,4-months) (n=6) and control BM cells (n=4). **(B)** Venn diagram analysis of tumor cells (n=3), yBM cells (1.5,4-months) (n=6) compared to the control BM cells (n=4). 303 are common genes between yBM and tumor cells. **(C)** Pathway enrichment analysis of differentially expressed genes between yBM and tumor cells. **(D)** PCA plot of BMT cells (n=3), yBM cells (1.5,4-months) (n=6) and control BM cells (n=4). **(E)** Venn diagram analysis of BMT cells (n=3), yBM cells (1.5,4-months) (n=6) compared to the control BM cells (n=4). 471 are common genes between yBM and BMT cells. **(F)** Pathway enrichment analysis of differentially expressed genes between yBM and BMT cells. **(G)** Gene Set Enrichment Analysis (GSEA) of differentially expressed genes between control and yBM/tumor cells. **(H)** GSEA of differentially expressed genes between control and yBM/BMT cells.

Altogether, these data imply that there is a high similarity between DKO yBM, BMT, and OS tumor genetic makeup, making yBM cells of DKO mice a possible surrogate for studying OS.

### *Trp53* SKO yBM is not tumorigenic

To test whether this phenotype is WWOX-dependent we generated a traceable mouse model harboring a*Trp53* KO alone and expressing tdTomato (SKO^TOM^) under the control of the same Osx1-Cre model (Figure 4S-A). SKO^TOM^ was validated using genomic PCR (Figure 4S-B). Importantly, SKO^TOM^ mice developed OS tumors at a median age of 11 months (data not shown), as previously shown(11).

We next validated the expression of our genetic manipulation in the BM cells of SKO^TOM^ mice (Figure 4S-C). Consistent with the DKO model, we observed an age-dependent increase in the percentage of tomato-positive cells (Figure 4S-D). In addition, we found that these tomato-positive BM cells were of mesenchymal origin (Figure 4S-E and F), like those observed in DKO^TOM^ mice. To further investigate the tumorigenic ability of SKO yBM cells, we injected them into NOD/SCID mice. None of the mice injected with SKO yBM cells formed tumors, even after 3-4 months of injection compared to DKO yBM cells where about 50% of the mice developed OS tumors (Figure 5S-A, B).

### WWOX loss results in Myc alteration in yBM cells

To uncover whether Myc is a unique event in DKO yBM cells, we subjected SKO yBM cells to RNA-seq analysis. We found low expression of Myc and its targets in SKO yBM as compared to those in DKO cells. In fact, the Myc target pathway was unique to yBM DKO cells (Figure 6A, B). Both RNA-seq and qPCR data analyses indicated enhanced upregulation of Myc targets in DKO cells relative to that in SKO yBM cells (Figure 6C, D). Further comparisons of SKO yBM with DKO tumor and BMT cells indicated that Myc targets are unique to DKO tumors and BMT cells (Figure 3S-C, D). Moreover, a comparison of SKO yBM cells and SKO tumor cells showed that Myc targets were unique to SKO tumor cells (Figure 3S-E). Importantly, Myc targets were mutually expressed in SKO and DKO tumors (Figure 3S-F), suggesting that the Myc pathway is indispensable at the tumor stage. These data indicate that Myc is a prominent pathway contributing to earlier stages of osteosarcomagenesis.

**Figure 6:**
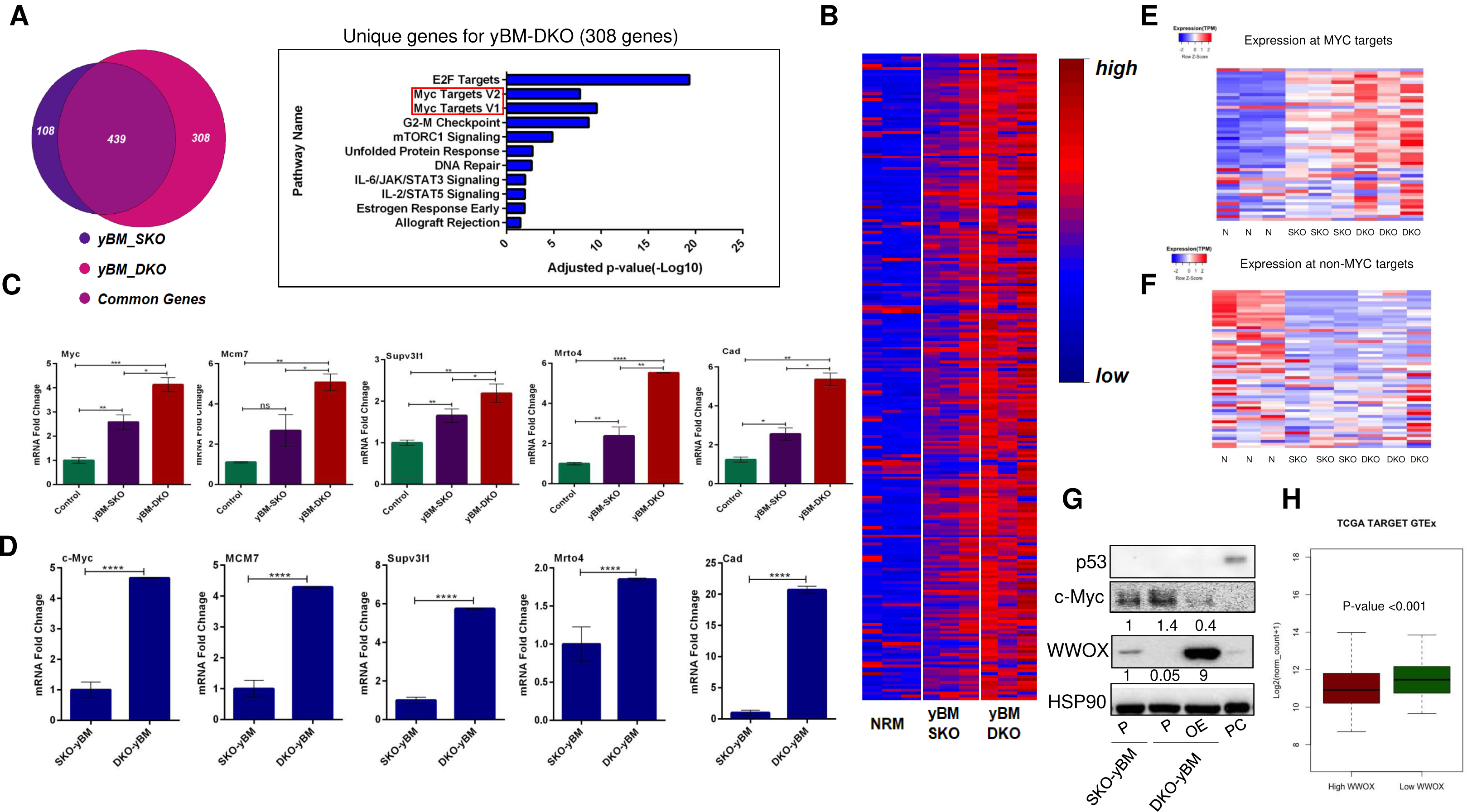
WWOX expression is inversely correlated with c-Myc in OS. **(A)** Venn diagram analysis of SKO and DKO yBM cells-left panel (n=3), 308 genes are unique for yBM DKO. Bar graph representing the significant pathways (Myc Targets) unique for DKO compared to SKO yBM-right panel. **(B)** Heat map illustration showing the differentially expressed genes between control BM (NRM), SKO and DKO yBM cells (n=3). **(C)** Bar graph-representing reads from RNA sequencing data for some Myc target genes in control BM, SKO and DKO yBM cells. **(D)** mRNA levels of Myc target genes in DKO and SKO yBM cells. **(E)** Color coded heatmap of normalized expression (TPM) of top 50 genes harbouring the highest Myc enrichment at promoters. **(F)** Color coded heatmap of normalized expression (TPM) of 50 genes depleted in Myc at promoters. **(G)** Immunoblotting of WWOX, c-Myc and p53 of SKO yBM and DKO yBM cells. P-Parental cell line, OE: Over expression, PC: positive control, HepG2 cells treated with nutlin (MDM2-inhibitor). Quantification of WWOX and c-Myc protein levels relative to SKO are shown. **(H)** Box Plot representing the correlation between WWOX and Myc expression in human osteosarcoma samples from TCGA TARGET GTEx. High WWOX represents samples >75^th^ percentile low WWOX represents <25^th^ percentile, y-axis represents Myc expression. PV: P-value.

To further confirm that Myc targets are certainly upregulated in DKO yBM cells, we performed Myc-chromatin immunoprecipitation and sequencing (ChIP-Seq) on DKO yBM cells (Figure 5S-G). We found that genes which are enriched in Myc at their promoters are indeed more expressed in the DKO compared to SKO yBM cells as opposed to genes depleted in Myc (Figure 6E, F 5S C-E). This data implies a regulatory role of WWOX on Myc targets expression in yBM cells. We further examined consequences of WWOX re-expression on Myc levels in DKO yBM cells and found a significant reduction in Myc levels upon WWOX restoration (Figure 6D). Remarkably, bioinformatical analysis of WWOX and Myc expression in human OS samples revealed a significant increase in Myc expression in human samples with low WWOX expression (Figure 6H). These findings further strengthen our hypothesis that depletion of WWOX results in Myc upregulation as an early event in osteosarcomagenesis.

### MCM7 upregulation in DKO yBM cells contributes to their tumorigenic ability

Molecular analysis revealed that the Myc pathway was one of the most prominent and significant early events in DKO yBM cells. This is consistent with previous studies demonstrating the upregulation of Myc in human OS(42),(43). Intriguingly, RNA-seq analysis also showed the upregulation of Myc targets (Figure 3S-G, H), including MCM7, which is related to cell cycle regulation and DNA replication(20). MCM7 upregulation was validated at the mRNA (Figure 7A) and protein levels as assessed by western blotting (Figure 7B-upper panel). We further detected a higher level of Myc and MCM7 in DKO yBM cells relative to that in SKO yBM cells (Figure 5S-F), which likely explains the higher survival and proliferative ability of DKO yBM cells (Figure 5S-G).

**Figure 7:**
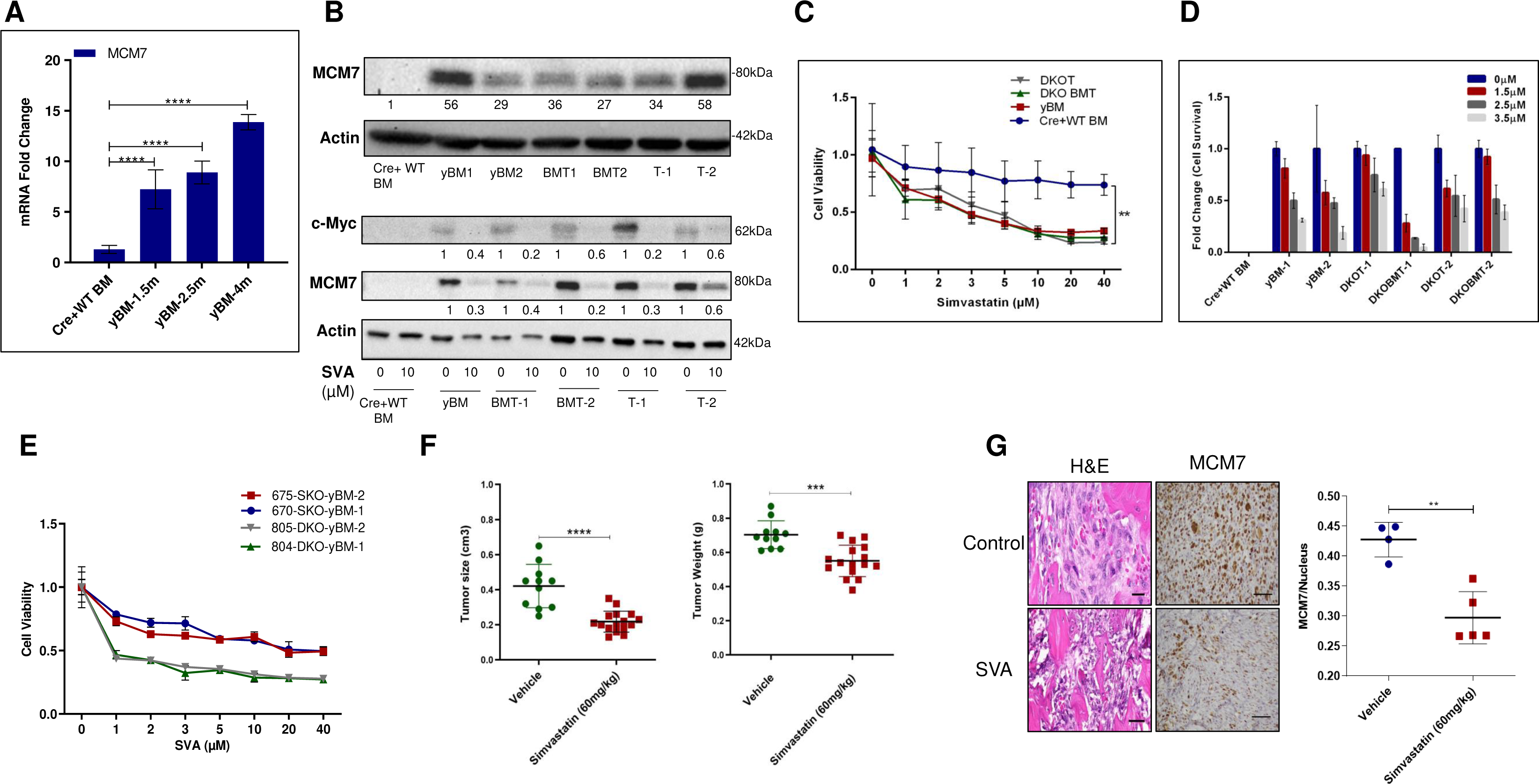
MCM7 upregulation promotes tumor progression. **(A)** mRNA levels of MCM7 in Cre+WT BM (Control), DKO yBM collected form 1.5, 2.5- and 4-months non-tumor bearing mice (n=3). ****p-value<0.0001. **(B)** Immunoblotting of MCM7 in Cre+WT BM (Control), DKO yBM, BMT and tumor cells (numbers indicate biological repeats). Quantification of MCM7 protein levels relative to Cre+ WT BM cells (Control) are shown-upper panel. Immunoblotting of MCM7 and c-Myc in Cre+WT BM (Control), DKO yBM, BMT and tumor cells (numbers indicate biological repeats) after SVA (µM) treatment for 24 hours-lower panel. Quantification of MCM7 and c-Myc protein levels relative to non-treated cells are shown. **(C)** MTT-proliferation assay of Cre+WT BM (Control), DKO yBM, BMT and tumor cells after SVA (µM) treatment for 24 hours. The graph represents fold change of cell viability relative to non-treated cells of each group. **(D)** Clonogenic assay of Cre+WT BM (Control), DKO yBM, BMT and tumor cells (numbers indicate biological repeats) after SVA (µM) treatment for 24 hours. The graph represents fold change of cell survival relative to non-treated cells of each group. **(E)** MTT-proliferation assay of SKO and DKO yBM cells (two biological repeats for each group) after SVA (µM) treatment for 24 hours. The graph represents fold change of cell viability relative to non-treated cells of each group. **(F)** Average size (cm^3^) and weight (g) of the vehicle or SVA (60mg/kg) treated tumors that developed in NOD/SCID mice after IT injection of yBM cells (* p-value <0.05, **p-value <0.01). **(G)** H&E staining of the Control (vehicle) and SVA treated tumors demonstrating the osteoblastic/fibroblastic characteristics of OS, IHC staining of MCM7 of the control (vehicle) and SVA treated tumors-left panel, quantification of the MCM7 positively stained nuclei compared to the control-right panel (**p-value<0.001).

Targeting MCM7 by statins has recently been shown to have an inhibitory effect on the proliferation of tumor cells(24). We next examined the effect of MCM7 inhibition on the proliferation of DKO yBM, BMT, and mouse OS tumor cells. We found that increased doses of simvastatin (SVA), previously shown to inhibit MCM7(24), inhibited the growth of yBM cells similar to that of OS tumor cells and BMT, but did not affect control BM cells (Figure 7C). In addition, we observed a dose-dependent inhibitory effect on the survival of DKO yBM, BMT, and tumor cells (Figure 7D). This treatment was accompanied by decreased MCM7 and Myc protein levels (Figure 7B-lower panel), consistent with a previous work(25). Moreover, the DKO yBM was more sensitive to SVA than the SKO yBM (Figure 7E).

Next, we examined the inhibitory effect of SVA on tumor growth *in vivo*. After tumor detection due to IT injection of DKO yBM in immunocompromised NOD/SCID mice, the mice were treated with vehicle or SVA (60 mg/kg/day) via gavage for an additional 10-days. We observed a significant decrease in the tumor size and weight following SVA treatment (Figure 7F). Interestingly, we also noticed a significant reduction in MCM7 levels in SVA-treated tumors, as validated by IHC (Figure 7G). Collectively, these data imply that MCM7, a Myc target gene, plays an important role in the early stages of osteosarcomagenesis.

## Discussion

High genetic heterogeneity and chromosomal instability contributes to worse prognosis(3). Hence, understanding the early molecular events in osteosarcomagenesis is crucial and could provide insights into early biomarkers and aid in the development of alternative therapies. In the present study, using the *Osterix1-cre* transgenic mouse model, we showed that deletion of *Wwox* and *Trp53* (DKO), while expressing the tdTomato reporter, resulted in the presence of genetically manipulated osterix-expressing cells in the BM. Interestingly, these genetically manipulated cells collected at the time of tumor detection expressed mesenchymal and osteoblastic markers, similar to OS tumor cells. In addition, BM cells isolated from DKO non-OS-bearing mice exhibited tumorigenic traits, as well as cellular and molecular changes resembling those of OS tumor cells and paired BM cells from OS-bearing mice. Our traceable OS mouse model facilitated examination and validation of molecular changes that occur during the early stages of osteosarcomagenesis. We found that Myc and its target genes were key players in the early stages of OS development. Inhibition of these targets may hamper the development of OS.

The cell of origin for OS remains controversial, with some studies suggesting that MSCs are the cell of origin(44). p53 has been extensively shown to play a central role in OS development in both human and mouse models(45),(28). Interestingly, p53 knockdown alone is not enough to transform human MSCs (hMSCs) to induce osteosarcomagenesis(44), which may indicate that at this stage, there is a need for additional hits to trigger the initiation of osteosarcoma. Consistent with these results, our data demonstrate that murine yBM cells collected from mice harboring the deletion of *Trp53* alone (SKO) were not tumorigenic, whereas yBM cells collected from DKO mice harboring the deletion of both *Wwox* and *Trp53* exhibited tumorigenic traits and closely resembled OS tumor cells. These findings indicate that loss of WWOX at earlier stages of tumorigenesis promotes osteosarcomagenesis. Our observation is in agreement with a previous study(44) that demonstrated that human MSCs transformed by Rb knockdown and c-Myc overexpression resulted in OS development and that these cells lost their mesenchymal stem cell markers and became more committed to the osteoblast lineage(44). Interestingly, we detected both mesenchymal and osteoblast markers in BM-MSCs using our yBM model. This may indicate that the specific genetic manipulation of the combined loss of WWOX and p53 under the *Osterix* promoter in our model could provide cells with plasticity to express both mesenchymal and osteoblast markers, thus transdifferentiating and contributing to the formation of a histological spectrum of OS.

OS is generally associated with alterations in the expression of genes involved in cell cycle regulation and apoptosis(46). Myc also plays an important role in the development and progression of OS. Notably, more than 10% of OS patients have c-Myc amplification(42),(43) and many more exhibit overexpression of Myc(15),(47). Interestingly, our yBM collected from a tumor free mouse and BMT cells share several genes with OS tumors. These genes are related to the cell cycle pathway, DNA repair, E2F and c-Myc targets. Detecting high levels of Myc in OS tumors is consistent with previous studies(15),(47), however its expression in yBM of DKO cells further strengthen that these cells resemble OS tumors and their indispensable role in osteosarcomagenesis.

Remarkably, our DKO yBM cells expressed high levels of Myc in comparison to SKO-yBM cells, and this differential expression of Myc and its target genes resulted in cellular advantage to DKO yBM, both *in vitro* and *in vivo*. Although c-Myc upregulation has been shown in p53-deficent OS tumors(48), our study provides clear evidence that combined deletion of WWOX and p53 resulted in increased Myc levels and activity in BM-MSCs as well as formation of indistinguishable OS tumors. This combined genetic manipulation seems to be critical for OS initiation since p53-deficient BM-MSCs, at the same stage, didn’t result in tumor formation in vivo further implying the significance of WWOX deletion in initiation of p53-deficient OS tumors.

A recent study in our lab demonstrated that WWOX suppresses Myc levels and activity in triple-negative breast cancer cell lines, by which WWOX binds the Myc transcription factor on the chromatin and modulates its activity on target genes. This negative regulation antagonizes epithelial-to-mesenchymal transition (EMT) and metastases(30). Moreover, in hepatocellular carcinoma (HCC) mouse model harboring deletion of Wwox in hepatocytes generated recently in our lab, it was shown that WWOX partially suppresses HCC through continuous suppression of Myc(49). These observations suggest an important role of tumor suppressor WWOX as a negative regulator of Myc in several cancer types. Nevertheless, this interaction between Myc and WWOX is largely unknown in OS. In our study, we show that genes that are enriched in Myc at their promoters are indeed more expressed in the DKO compared to SKO yBM as validated by Myc-ChIP-seq. Moreover, WWOX restoration to DKO yBM cells negatively regulates Myc protein levels. Moreover, bioinformatical analysis of human OS samples from TCGA demonstrates the inverse correlation between WWOX and Myc. This data further strengthens the regulatory role of WWOX on Myc in OS.

Given that c-Myc has been shown to cooperate with other tumor suppressor genes to transform hMSCs to form OS(44), we assumed that our model recapitulates human OS and that the combined deletion of WWOX and p53 is critical for the initiation of OS. Importantly, we propose that yBM cells can be used as surrogates to investigate the early stages of osteosarcomagenesis. Moreover, our study validates that the BM-MSCs with co-deletion of WWOX and p53 are tumorigenic compared to p53 deletion alone although with a later latency compared with BM-MSCs with p53 and Rb co-deletion(50). Furthermore, our DKO yBM model exhibited early molecular pathway enrichment, which is consistent with a recent multi-omics analysis performed on human OS clinical samples(51). Interestingly, Myc amplification and overexpression of Myc targets were detected and correlated with a worse prognosis, indicating the crucial role of Myc in osteosarcomagenesis. Moreover, this analysis of human OS clinical samples further indicated the alteration of several tumor suppressor genes, including *TP53* and *WWOX* deletion, which further strengthens our model recapitulates human OS. In addition, it also showed that deletion of *RB1*, resulting in the attenuation of cell cycle-related pathways, which is also one of the most prominently altered pathways in our model. Furthermore, our DKO model showed an upregulation of E2F targets, which is known to be correlated with OS worse prognosis(52). Moreover, a recent study indicated a role of Myc in controlling the activation of E2F genes and the possibility that deregulation of Myc contributes in multiple ways to the activation of the E2F/Rb pathway(53). This further indicates the critical role of Myc in OS development and progression. Of note, a single-cell RNA sequencing performed on human OS samples and the construction of a novel OS classification model revealed three subtypes of OS that exhibit distinct prognoses and different expression levels of therapy-related genes(54), where some of these genes are differentially expressed in murine DKO-yBM cells, such as Myc and CDK4.

In addition to the role of Myc itself, we validated Myc activity by examining one of the c-Myc targets, MCM7. MCM7 is part of the MCM complex, that plays an important role in cell cycle initiation, and is upregulated in several tumors, including sarcomas(21),(22). Interestingly, DKO yBM showed upregulation of MCM7 in an age-dependent manner at both mRNA and protein levels. High levels of MCM7 protect the cell by acting as a guardian and inducing genomic stability, thus making the cells more resistant to replication stress caused by chemotherapy(24). As MCM complex components are present in excess in the cells and act as a dormant origin, once these excess amounts are depleted, the cells will be more sensitive to replicative stress(23), making them a good therapeutic target.

Statins are competitive inhibitors of 3-hydroxy-3-methylglutaryl-coenzyme A (HMG-CoA) reductase, a rate-limiting enzyme that converts HMG-CoA to mevalonate during cholesterol synthesis(55). In addition to their traditional role in lowering serum cholesterol levels, extensive studies have suggested that statins inhibit carcinogenesis(56),(57),(58) and can be used in cancer therapy(59),(60). Furthermore, studies suggested that statin administration prior to cancer diagnosis exhibited reduced cancer related mortality by up to 15%(61) and showed that long term use of statins reduces the risk of colorectal cancer(62).

Interestingly, MCM7 can be targeted by statins, generating more DNA damage and inhibiting cell proliferation(63),(25). Our data demonstrate that MCM7 inhibition by SVA reduced cell proliferation and survival in a dose-dependent manner. Importantly, DKO yBM, but not SKO yBM, showed less survival ability and exhibited a higher sensitivity. The significant inhibitory effect of SVA can be attributed to the central role of MCM7 in cell fate. It is also possible that SVA has effects that go beyond inhibition of MCM7. In addition, SVA treatment *in vivo* resulted in a reduction in tumor size and weight, which was accompanied by a decrease in MCM7 protein levels.

## Conclusions

Consistent with their roles as tumor suppressor genes, the combined deletion of WWOX and p53 in Osx1-positive progenitor cells resulted in their transformation and growth advantage. This advantage is mediated, at least in part, by Myc overexpression, likely during early stages of OS cell proliferation and survival. Given that p53 alone is not sufficient to drive the early molecular changes observed, WWOX silencing and Myc overexpression may be considered targets for the prevention and treatment of OS patients. Our study provides a cellular and genetic platform to study the molecular mechanisms underlying OS transformation, progression, and intervention.

## Acknowledgments

We are grateful to Dr. Sri Repudi, Ms. Natalie Kaufman and Randa Teswak for their help and technical assistance. The Aqeilan’s lab is funded by the European Research Council (ERC) [No. 682118] to R.I.A and the Bi-national Science Foundation (BSF) [No. 2017269] to R.I.A and G.S.S. Osama Hidmi is in part supported by The Carole and Andrew Harper Diversity PhD Program.

## Author Contributions

Conceptualization: RA and RIA; Methodology: RA, OH, AH-Y, and JM; Investigation: RA, OH, AH-Y, JM, and JD; Writing—review and editing: RA and RIA; Funding acquisition: RIA; Resources: RA, OH, and AH-Y; Project administration and supervision; RIA.

