## Supplemental Figures and Table for "WWOX promotes osteosarcoma development via upregulation of Myc"

### Supplementary Figure Legends

#### ***Supplementary Figure 1: Immunophenotyping of BMT and tumor cells reveals mesenchymal/osteoblastic identity.***

**(A)** Flow cytometry analysis (histogram) of directly collected and/or cultured BM from a Cre- mouse (Cre-Direct BM), tumor (Direct DKOT/Cult DKOT) and paired BM cells (Direct DKO BMT/Cult DKO BMT) collected from DKO mouse demonstrating the expression of mesenchymal (Sca-1 and CD29) vs. hematopoietic (CD45.2) markers in tdTomato positive cells; representative experiment. **(Bi, Bii)** A bar graph demonstrating the percentage of hematopoietic and mesenchymal population in direct and cultured BMT and tumor cells respectively (n=3). **(C)** mRNA levels of osteoblast markers in DKO BMT and tumor (DKOT) cells vs WT BM (Cre+ WT BM) cells (n=3) (**p-value<0.01, ***p-value <0.001, ****p-value<0.0001, ns: not significant). **(D)** Alizarin Red Staining demonstrating calcium deposition during osteoblast differentiation of tumor, BMT and yBM cells. **(E)** mRNA levels of osteoblast differentiation markers of tumor, BMT and yBM cells. **US**: Unstained, **S**: Stained, **Direct**: Directly collected BM before culturing, **Cult**: Cultured.

#### ***Supplementary Figure 2: young BM (yBM)-derived cells are of mesenchymal/osteoblastic identity and exhibit metastatic ability.***

**(A)** Flow cytometry analysis (histogram) of directly collected and/or cultured BM from a Cre negative mouse (Cre-BM-2m), young BM (DKO-Direct BM-2m) demonstrating the expression of mesenchymal (Sca-1 and CD29) vs. hematopoietic (CD45.2) markers in tdTomato positive cells; representative experiment. **(B)** A bar graph demonstrating the percentage of hematopoietic and mesenchymal population in direct and cultured yBM cells (n=3). **(C)** mRNA levels of osteoblast markers in yBM and WT BM cells (Cre+WT BM). (** p-value< 0.01, ****p-value<0.0001). **(D)** Wound healing assay-migration. Time plot representing the Relative Wound Density of yBM, BMT and tumor cells (n=3). **(F)** Wound healing assay-invasion. Time plot representing the Relative Wound Density of yBM, BMT and tumor cells (n=3). **US**: Unstained, **S**: Stained, **Direct**: Directly collected BM before culturing, **Cult**: Cultured.

***Supplementary Figure 3: Early Genes Contributing to Osteosarcomagenesis.***

**(A)** Heat map of the 303 mutual genes between yBM and tumor cells represented in the venn diagram in Fig 5B. **(B)** Heat map of the DE genes between yBM (n=6) and BMT (n=3) compared to the control BM cells (n=4). **(C)** Venn diagram analysis of yBM SKO and Tumor DKO (Tum-DKO) representing the common and unique genes between them-upper panel, Bar graph representing the significant pathways (Myc Targets) unique for DKO tumor cells compared to SKO yBM cells-lower panel. **(D)** Venn diagram analysis of yBM SKO and BMT DKO, representing the common and unique genes between them-upper panel, Bar graph representing the significant pathways (Myc Targets) unique for DKO BMT cells compared to SKO yBM cells-lower panel. **(E)** Venn diagram analysis of SKO yBM and SKO tumor cells representing the common and unique genes between them-upper panel, Bar graph representing the significant pathways (Myc Targets) unique for SKO tumor cells compared to SKO yBM cells-lower panel. **(F)** Venn diagram analysis of SKO and DKO tumor cells representing the common and unique genes between them-upper panel, Bar graph representing the significant pathways (Myc Targets) mutual between DKO and SKO tumor cells-lower panel. **(G)** Heat map of the 40 Myc target genes that are upregulated in yBM, and tumor cells compared to control cells. **(H)** Heat map of the 39 Myc target genes that are upregulated in yBM and BMT cells compared to control cells.

***Supplementary Figure 4: Generation of a traceable mouse model of SKO-p53 osteosarcoma cell.***

**(A)** Schematic illustration of the experimental SKO-p53 OS mouse model. **(B)** Genomic PCR of Cre negative (Cre-), Cre positive (Cre+ WT) and Single knock out ($SKO^{TOM}$) and Double knock out ($DKO^{TOM}$) mice. Labels in the left panel represent the primer pairs and in the right panel represent the expected band size. **(C)** Immunoblotting of WWOX, p53 and tdTomato in the SKO BM, DKO BM cells before (0 min) and after (30, 120 min) of ionizing radiation exposure (IR-10Gy). PC-(positive control)-HEPG2 cells treated with nutlin (MDM2-inhibitor). **(D)** Summary graph of flow cytometry analysis of the percentage of tdTomato positive cells in DKO BM, SKO BM and wild type BM cells (Cre+WT BM) at different ages. **(E)** A bar graph demonstrating the percentage of hematopoietic and mesenchymal population in direct and cultured yBM SKO cells (n=3). **(F)** Flow cytometry analysis (histogram) of directly collected and cultured BM from a yBM-SKO demonstrating the expression of mesenchymal (Sca-1 and CD29) vs. hematopoietic (CD45.2) markers in tdTomato positive cells; representative experiment. **WT**: Wildtype, **SKO**: Single Knock Out, **DKO**: Double Knock Out, **BM**: Bone Marrow.

***<u>Supplementary Figure 5: MCM7 upregulation in DKO yBM cells promotes their tumorigenicity.</u>***

**(A)** Representative images of the OS tumors developed in NOD/SCID mice after intratibial (IT) injection of DKO-yBM cells compared to SKO-yBM-upper panel, H&E of the developed tumors-lower panel. Histological validation indicates OS characteristics. **(B)** Bar graph representing the percentage of immunocompromised mice that developed OS after IT injection of SKO and DKO yBM cells. **(C)** Color coded heatmap of normalized expression (TPM) of top 100 genes harbouring the highest Myc enrichment at promoters. (D) Color coded heatmap of normalized expression (TPM) of 100 genes depleted in Myc at promoters. **(E)** Gene snapshots of examples of the MYC-enriched promoters. **(F)** Immunoblotting of MCM7 and c-Myc in SKO-yBM and DKO-yBM cells (the numbers indicate biological repeats), quantification of MCM7 and Myc protein levels relative to SKO yBM cells are shown. **(G)** Quantification of colony formation of SKO-yBM and DKO-yBM (n=3). ****p-value<0.0001.

***<u>Supplemental Tabel 1: qPCR primers list and their sequences along with the predicted sizes.</u>***

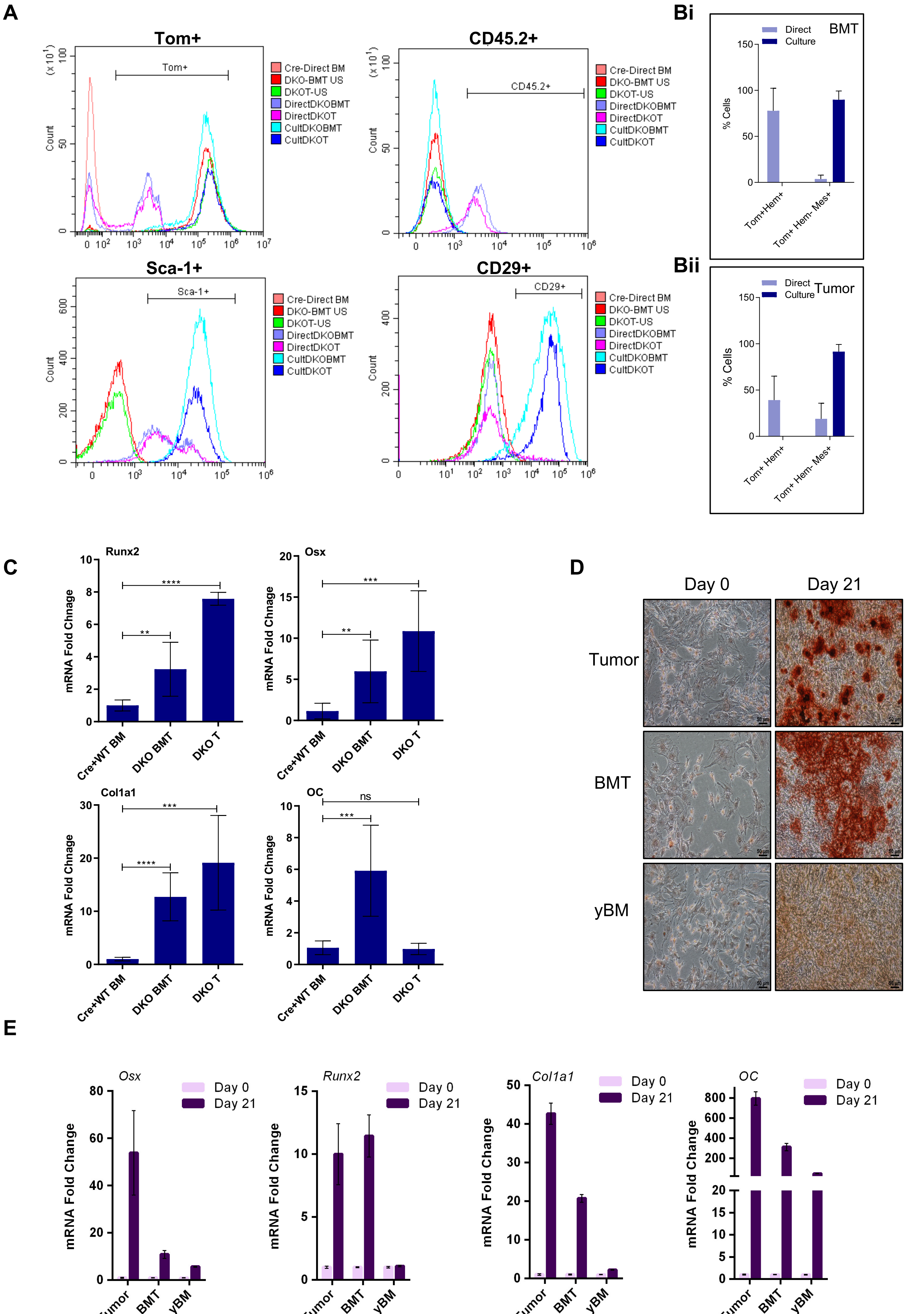

A
Tom+
Tom+
CD45.2+
CD45.2+
Sca-1+
Sca-1+
CD29+
CD29+
Count
Cre-Direct BM
DKO-BMT US
DKOT-US
DirectDKOBMT
DirectDKOT
CultDKOBMT
CultDKOT
Bi
BMT
Direct
Culture
% Cells
Tom+Hem+
Tom+ Hem- Mes+
Bii
Tumor
C
Runx2
Osx
Col1a1
OC
mRNA Fold Chnage
Cre+WT BM
DKO BMT
DKO T
ns
D
Day 0
Day 21
Tumor
BMT
yBM
E
Osx
Runx2
Col1a1
OC
mRNA Fold Change
Day 0
Day 21
Tumor
BMT
yBM


Figure 1S

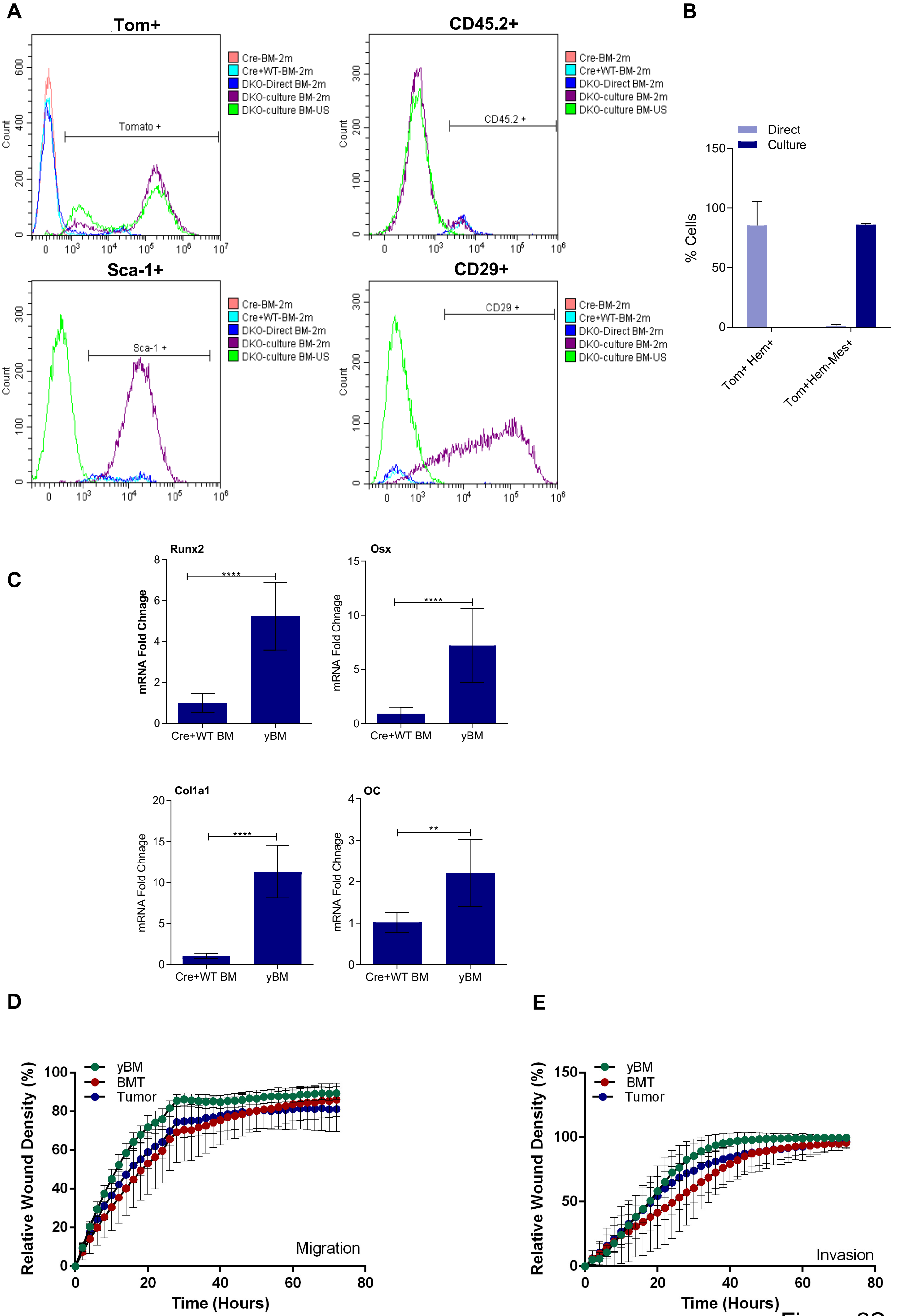
A
Tom+
Tomato +
Count
CD45.2+
CD45.2 +
Sca-1+
Sca-1 +
CD29+
CD29 +
Cre-BM-2m
Cre+WT-BM-2m
DKO-Direct BM-2m
DKO-culture BM-2m
DKO-culture BM-US
B
Direct
Culture
% Cells
Tom+ Hem+
Tom+Hem-Mes+
C
Runx2
Osx
Col1a1
OC
mRNA Fold Chnage
Cre+WT BM
yBM
****
**
D
E
yBM
BMT
Tumor
Relative Wound Density (%)
Time (Hours)
Migration
Invasion

Figure 2S

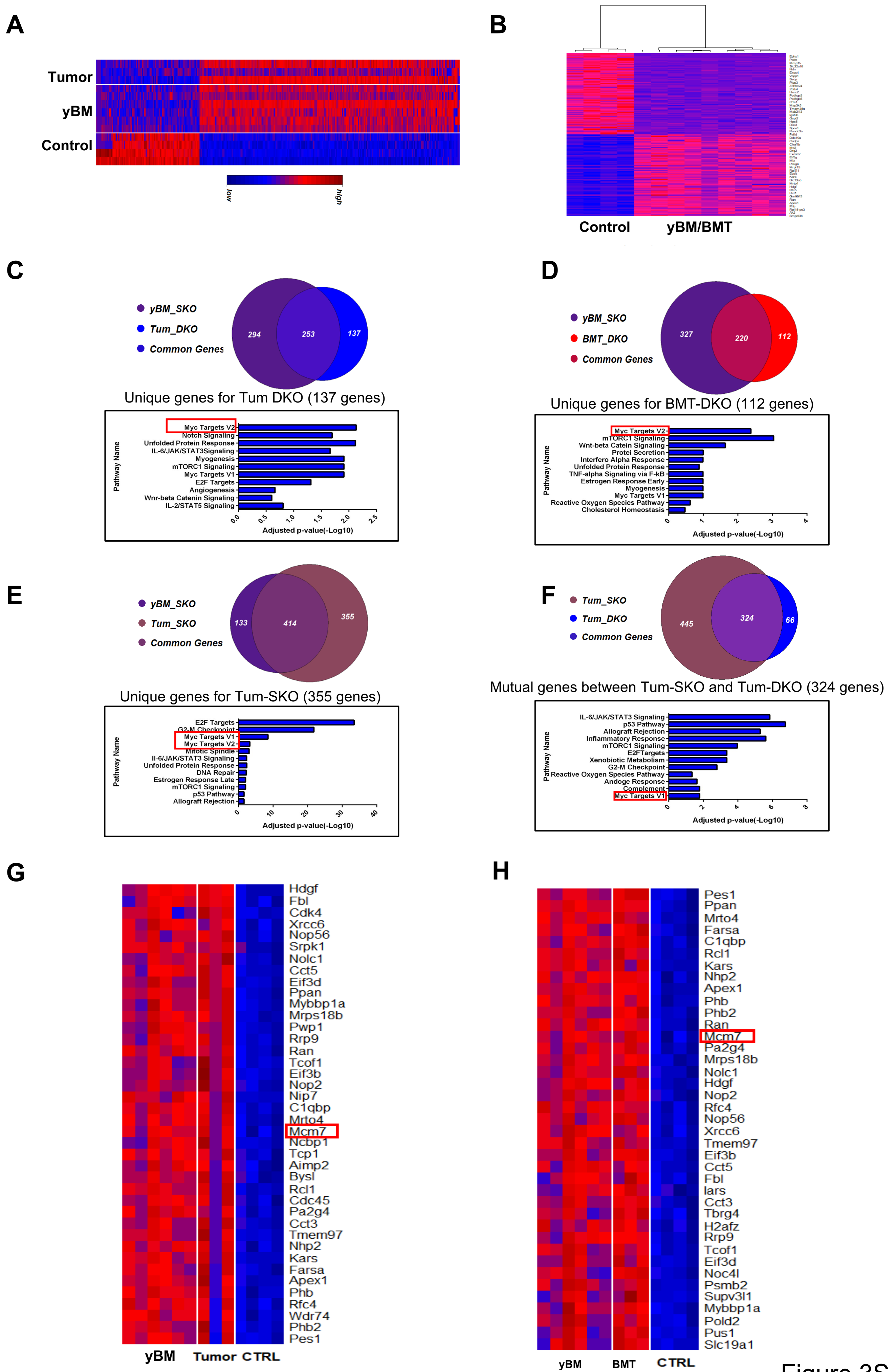

A
Tumor
yBM
Control
low
high
B
Control
yBM/BMT
C
yBM_SKO
Tum_DKO
Common Genes
294
253
137
Unique genes for Tum DKO (137 genes)
Pathway Name
Myc Targets V2
Notch Signaling
Unfolded Protein Response
IL-6/JAK/STAT3Signaling
Myogenesis
mTORC1 Signaling
Myc Targets V1
E2F Targets
Angiogenesis
Wnr-beta Catenin Signaling
IL-2/STAT5 Signaling
0.0 0.5 1.0 1.5 2.0 2.5
Adjusted p-value(-Log10)
D
yBM_SKO
BMT_DKO
Common Genes
327
220
112
Unique genes for BMT-DKO (112 genes)
Pathway Name
Myc Targets V2
mTORC1 Signaling
Wnt-beta Catein Signaling
Protei Secretion
Interfero Alpha Response
Unfolded Protein Response
TNF-alpha Signaling via F-kB
Estrogen Response Early
Myogenesis
Myc Targets V1
Reactive Oxygen Species Pathway
Cholesterol Homeostasis
0 1 2 3 4
Adjusted p-value(-Log10)
E
yBM_SKO
Tum_SKO
Common Genes
133
414
355
Unique genes for Tum-SKO (355 genes)
Pathway Name
E2F Targets
G2-M Checkpoint
Myc Targets V1
Myc Targets V2
Mitotic Spindle
Il-6/JAK/STAT3 Signaling
Unfolded Protein Response
DNA Repair
Estrogen Response Late
mTORC1 Signaling
p53 Pathway
Allograft Rejection
0 10 20 30 40
Adjusted p-value(-Log10)
F
Tum_SKO
Tum_DKO
Common Genes
445
324
66
Mutual genes between Tum-SKO and Tum-DKO (324 genes)
Pathway Name
IL-6/JAK/STAT3 Signaling
p53 Pathway
Allograft Rejection
Inflammatory Response
mTORC1 Signaling
E2FTargets
Xenobiotic Metabolism
G2-M Checkpoint
Reactive Oxygen Species Pathway
Andoge Response
Complement
Myc Targets V1
0 2 4 6 8
Adjusted p-value(-Log10)
G
Hdgf
Fbl
Cdk4
Xrcc6
Nop56
Srpk1
Nolc1
Cct5
Eif3d
Ppan
Mybbp1a
Mrps18b
Pwp1
Rrp9
Ran
Tcof1
Eif3b
Nop2
Nip7
C1qbp
Mrto4
Mcm7
Ncbp1
Tcp1
Aimp2
Bysl
Rcl1
Cdc45
Pa2g4
Cct3
Tmem97
Nhp2
Kars
Farsa
Apex1
Phb
Rfc4
Wdr74
Phb2
Pes1
yBM
Tumor
CTRL
H
Pes1
Ppan
Mrto4
Farsa
C1qbp
Rcl1
Kars
Nhp2
Apex1
Phb
Phb2
Ran
Mcm7
Pa2g4
Mrps18b
Nolc1
Hdgf
Nop2
Rfc4
Nop56
Xrcc6
Tmem97
Eif3b
Cct5
Fbl
Iars
Cct3
Tbrg4
H2afz
Rrp9
Tcof1
Eif3d
Noc4l
Psmb2
Supv3l1
Mybbp1a
Pold2
Pus1
Slc19a1
yBM
BMT
CTRL


Figure 3S

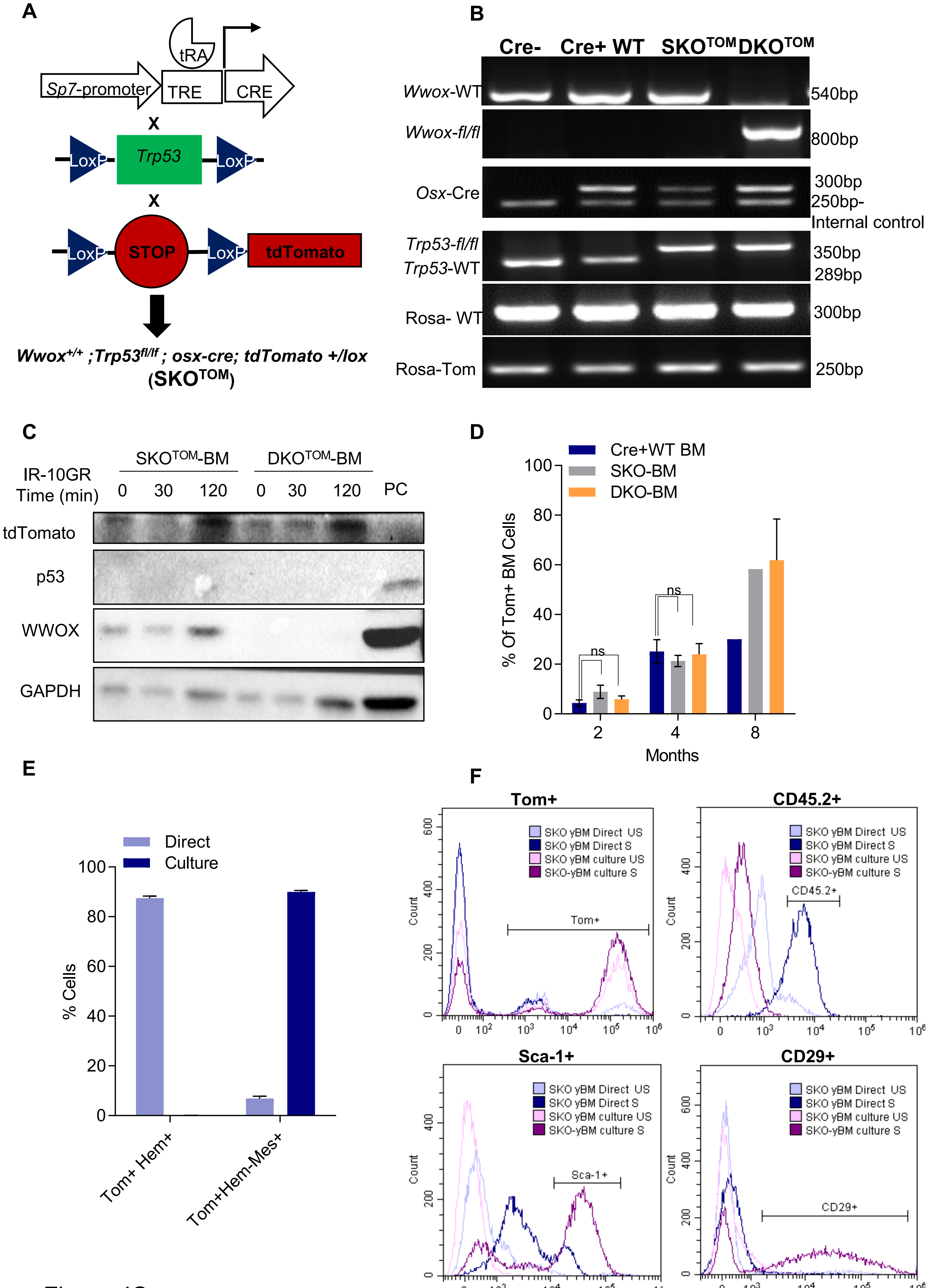

A
tRA
Sp7-promoter
TRE
CRE
X
LoxP
Trp53
LoxP
X
LoxP
STOP
LoxP
tdTomato
Wwox+/+ ;Trp53fl/lf ; osx-cre; tdTomato +/lox
(SKOTOM)
B
Cre-
Cre+ WT
SKOTOM
DKOTOM
Wwox-WT
540bp
Wwox-fl/fl
800bp
Osx-Cre
300bp
250bp-
Internal control
Trp53-fl/fl
Trp53-WT
350bp
289bp
Rosa- WT
300bp
Rosa-Tom
250bp
C
SKOTOM-BM
DKOTOM-BM
IR-10GR
Time (min)
0 30 120 0 30 120 PC
tdTomato
p53
WWOX
GAPDH
D
Cre+WT BM
SKO-BM
DKO-BM
% Of Tom+ BM Cells
ns
ns
Months
2 4 8
E
Direct
Culture
% Cells
Tom+ Hem+
Tom+Hem-Mes+
F
Tom+
CD45.2+
Sca-1+
CD29+
SKO yBM Direct US
SKO yBM Direct S
SKO yBM culture US
SKO-yBM culture S
Count
Tom+
CD45.2+
Sca-1+
CD29+


Figure 4S

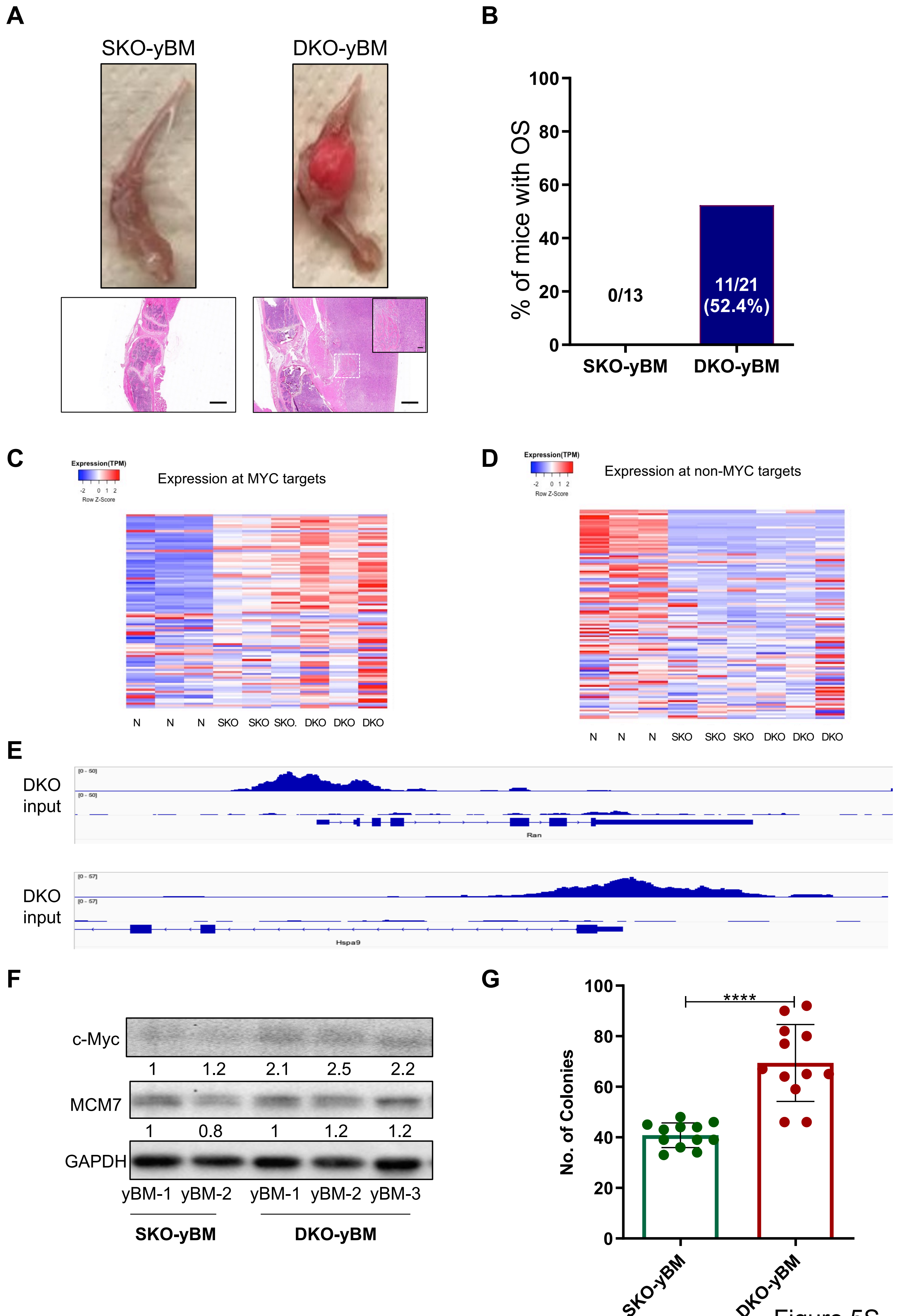
A
SKO-yBM
DKO-yBM
B
% of mice with OS
100
80
60
40
20
0
0/13
11/21
(52.4%)
SKO-yBM
DKO-yBM
C
Expression(TPM)
-2 0 1 2
Row Z-Score
Expression at MYC targets
N N N SKO SKO SKO. DKO DKO DKO
D
Expression(TPM)
-2 0 1 2
Row Z-Score
Expression at non-MYC targets
N N N SKO SKO SKO DKO DKO DKO
E
DKO
input
[0 - 50]
[0 - 50]
Ran
DKO
input
[0 - 57]
[0 - 57]
Hspa9
F
c-Myc
1 1.2 2.1 2.5 2.2
MCM7
1 0.8 1 1.2 1.2
GAPDH
yBM-1 yBM-2 yBM-1 yBM-2 yBM-3
SKO-yBM
DKO-yBM
G
No. of Colonies
100
80
60
40
20
0
****
SKO-yBM
DKO-yBM

Figure 5S

| Gene | | Sequence | Product size |
| --- | --- | --- | --- |
| Osterix | Osx | F:CCCTTCTCAAGCACCAATGG | |
| | | R:AGGGTGGGTAGTCATTTGCATAG | 85 bp |
| RUNX2 | RUNX2 | F:CGGCCCTCCCTGAACTCT | |
| | | R:TGCCTGCCTGGGATCTGTA | 75 bp |
| Colagen1A1 | Col1A1 | F:CTTGGTGGTTTTGTATTCGATGAC | |
| | | R:GCGAAGGCAACAGTCGCT | 101 bp |
| Osteocalcin | OC | F:CTGACAAAGCCTTCATGTCCAA | |
| | | R:GCGCCGGAGTCTGTTCACTA | 59 bp |
| MCM7 | MCM7 | F:AGTATGGGACCCAGTTGGTTC | |
| | | R:GCATTCTCGCAAATTGAGTCG | 115 bp |
| c-Myc | c-Myc | F:ATGCCCCTCAACGTGAACTTC | |
| | | R:CGCAACATAGGATGGAGAGCA | 228 bp |
| HPRT | HPRT | F:TCAGTCAACGGGGGACATAAA | |
| | | R:GGGGCTGTACTGCTTAACCAG | 142 bp |
| Supv3l1 | Supv3l1 | F:CTGGCAGATTCAGCTCACAC | |
| | | R:TGCCCATCAACTTGTGCAAA | 223 bp |
| Cad | Cad | F:GCAAGTGGTTTGAATCCTCGG | |
| | | R:CATTGGGGTCCACGAATGG | 246 bp |
| Mrto4 | Mrto4 | F:GCCAACATGAGGAACAGCAA | |
| | | R:AGCCCAACTTCACCTCTCAA | 170 bp |
